# A low-cost, open-source 3D printer for multimaterial and high-throughput direct ink writing of soft and living materials

**DOI:** 10.1101/2024.10.01.615991

**Authors:** Jonathan D. Weiss, Alana Mermin-Bunnell, Fredrik S. Solberg, Tony Tam, Luca Rosalia, Amit Sharir, Dominic Rütsche, Soham Sinha, Perry S. Choi, Masafumi Shibata, Yellappa Palagani, Riya Nilkant, Kiruthika Paulvannan, Michael Ma, Mark A. Skylar-Scott

**Author notes:** **Corresponding Author**, X (Twitter): @mascott85. These authors contributed equally (Jonathan D. Weiss, Alana Mermin-Bunnell).

## Abstract

Direct ink writing is a 3D printing method that is compatible with a wide range of structural, elastomeric, electronic, and living materials, and it continues to expand its uses into physics, engineering, and biology laboratories. However, the large footprint, closed hardware and software ecosystems, and expense of commercial systems often hamper widespread adoption. Here, we present a compact, simple-to-build, low-cost, multimaterial, and high-throughput direct ink writing 3D printer platform with detailed assembly files and instructions provided freely online. In contrast to existing low-cost 3D printers and bioprinters, which typically modify off-the-shelf plastic 3D printers, this system is built from scratch, offering a lower cost and full customizability. Despite its low cost, we demonstrate advanced active mixing and multimaterial multinozzle 3D (MM3D) printing methods, which previously have relied on expensive and custom motion control platforms. We finally introduce embedded multinozzle and 3D gradient nozzle designs that offer high throughput and graded 3D parts. This powerful, easy-to-build, and customizable printing platform can help stimulate a vibrant biomaker community of engineers, biologists, and educators.

## 1. Introduction

Direct ink writing (DIW) is a versatile and rapidly advancing method for 3D printing soft and biological viscoelastic materials.^[1]^ In contrast with other 3D printing methods such as inkjet and stereolithography (SLA), DIW is ideally suited to printing multiple materials and can work with a wide spectrum of ink viscosities and chemistries.^[2]^ Thus, DIW has found a broad range of applications from printed biological tissues,^[3–7]^ 3D soft robots,^[8,9]^ energy and electronic materials,^[10–13]^ and structural light-weight and metamaterial lattices.^[14–16]^ Complementing this diverse materials palette is a collection of different DIW modalities, including embedded 3D printing,^[3,17,18]^ multimaterial gradients,^[19]^ chaotic,^[20]^ core-shell,^[21]^ material switching,^[22]^ and multimaterial multinozzle 3D printing.^[9]^

However, existing commercial printers for DIW and 3D bioprinting remain expensive, typically ranging from $10,000-$200,000, and often present researchers with a closed ecosystem of unmodifiable hardware, materials, and software.^[23]^ We posit that open source and open access hardware is ideally suited for keeping pace with rapid advancements in 3D printing, allowing free modification of all components. However, existing open source 3D printers usually modify off-the-shelf x, y, z-gantry systems,^[23–32]^ robotic arms,^[33]^ or even compact disc drives^[34]^ (Table S1). Their frequent reliance on commercial backbones can add significant cost, reduce design versatility, and rely on continued support and production of the base hardware that is being modified. In contrast, a bottom-up 3D printer design can offer greater flexibility, allowing on-the-fly customization of printer build volume, motors and axes, and printheads.

Here, we present the “Printess,” a $250 bottom-up open-source 3D printer constructed from six linear actuators and drivers, a microcontroller, and 3D-printed and laser-cut components. The printer is lightweight (3kg), compact (23 cm x 23 cm x 40 cm), and can be carried with one hand into and out of a sterile environment such as a biosafety cabinet. Despite its low cost, the printer exhibits a high degree of motion accuracy, achieving 10-µm motion repeatability. Underscoring the versatility of the Printess, we demonstrate its use in multimaterial bioprinting of a wide variety of inks, curing chemistries, and applications, including cell-laden biological materials for tissue printing, photocurable hydrogels for trileaflet valve printing, and thermally curing elastomers for soft auxetic multimaterial printing. Most importantly, the Printess supports many emerging advanced printing modalities, including multimaterial, multimaterial active mixing, multimaterial multinozzle, and embedded 3D printing. Two of these methods, active mixing^[22]^ and multimaterial multinozzle 3D printing,^[9]^ were previously demonstrated using custom motion control stages costing over $300,000, hindering widespread adoption. To illustrate the versatility of the Printess, we further present two novel printing modalities: embedded active mixing and embedded multinozzle printing to produce gradients and high-throughput arrays of 3D geometries, respectively. To facilitate adoption and modification, a comprehensive list of drawings, instructions, code, and editable 3D files for the printer and its nozzles are provided online.

## 2. Results and Discussion

### 2.1. Design and Construction

The Printess chassis is constructed from two laser-cut acrylic sheets and 3D printed supports and adaptors (**Figure 1**). All components are screw-mounted using heat set inserts, allowing repeated assembly and disassembly from a boxed kit to facilitate transportation, customization for research, and use in hands-on student learning workshops. These structural components are used to mount six NEMA 11 linear actuators: base-mounted x- and y-axes (100 mm) that move the print bed, two mounted z-axes (100 mm) that can independently translate a pair of custom syringe pumps above, and two extrusion axes (50 mm) for volumetric material deposition. The collective build dimensions are 80mm x 80mm x 80mm. The six linear axes are controlled by an open source RepRap RUMBA+ 3D printer board housing six stepper motor drivers. The microcontroller operates on an ATmega2560 microcontroller running a six linear axis version of the Marlin computer numerical control (CNC) firmware. The Printess can also be expanded with a number of optional features, including fans, heat beds with temperature control, a touchscreen, and end-stops. The printer is connected via USB, can be operated through the Pronterface graphical user interface, and is compatible with machines running Windows or MacOS.

**Figure 1:**
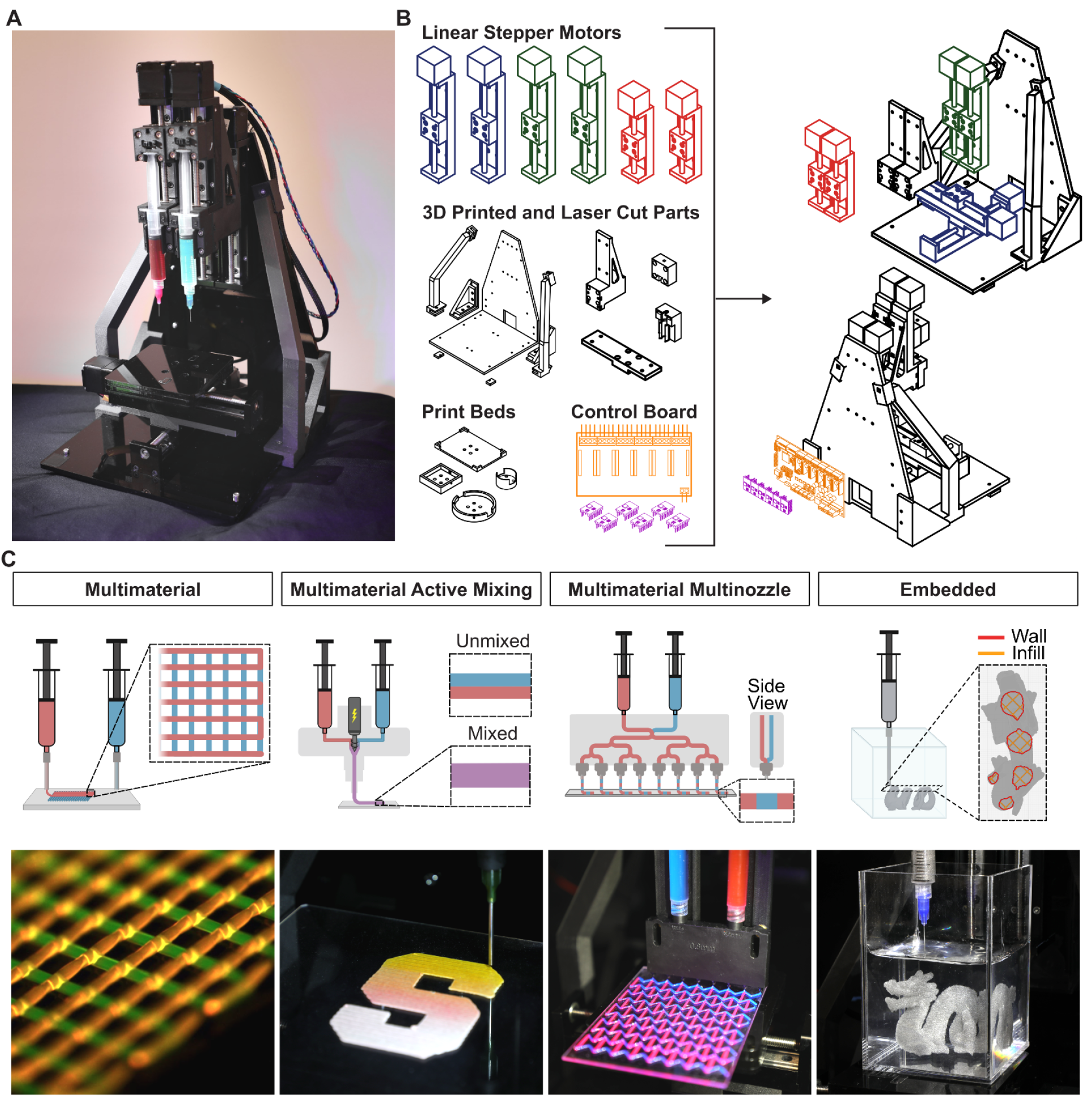
Design and Overview. **A)** Photograph of the assembled Printess with two mounted syringes. **B)** Overview of the Printess’ major components (left) and exploded front and back views (right). **C)** Overview of various direct ink writing modalities with photographs of representative prints in progress.

At its core, the six-axis machine is designed for multimaterial 3D printing of soft materials with two syringe pump extruders. The syringe pump can house 1, 3, 5, and 10 mL disposable and sterile syringes that are installed by a snap-fit geometry and can be temperature-controlled with a custom syringe cooling sleeve (Supporting Methods). This snap-fit design avoids having to sandwich the syringe barrel with a screw-on barrel holder and ensures an uninterrupted view of the contents of the syringe throughout printing for debugging, process monitoring, and educational purposes. The extruded material is deposited onto a print bed with a bed leveler and sample holder that can be customized for glass slides, Petri dishes, multiwell plates, or a temperature-controlled metal plate (Supporting Methods). In addition to these core components for multimaterial and embedded 3D printing, the build can be modified with additional components to enable active gradient multimaterial mixing and multimaterial multinozzle 3D printing.

A complete parts list for the Printess can be found in the Supporting Information with all purchased components and raw materials costing approximately $250 (Table S2). We have tested manufacturing the Printess in two separate laboratories using printed components from two different commercial fused deposition modeling (FDM) 3D printers loaded with two different filament materials: 1) Prusa I3 MK3S loaded with poly(lactic acid) (PLA) and 2) Markforged Mark II printer loaded with nylon with chopped carbon fibers (Onyx) inlaid with continuous carbon fibers (Figure S1). Note that depending on 3D printer performance and settings, the 3D printed component dimensions may need to be adjusted; editable 3D files are provided for this purpose. Once components are printed and laser cut, the printer can be assembled in under one hour. To date, our laboratory has constructed a fleet of sixteen Printesses (Figure S2), where they find daily use for research, education, and workshops.

### 2.2. Characterization and Multimaterial Printing

The Printess linear actuators are driven by NEMA 11 stepper motors. To reduce printer cost and complexity, the Printess does not use encoders, so the actuators are driven in an open-loop fashion. To measure the error associated with stepper motor skipping or thread backlash, we commanded a 10 mm sawtooth motion in the x-, y-, or z-axes while measuring true linear position using an external optical linear encoder. Both the z-axes were commanded as a dual carriage. The resulting error between real and nominal distances as the motors change direction between the forward and backward directions arises chiefly from thread backlash, which is about ±75 µm for the x-axis, ±150 µm for the y-axis, and ±50 for the z-axes (**Figure 2A-B)**. These backlash errors are similar to other published bioprinter designs,^[25]^ and can be reduced by compensatory motion to overcome thread backlash upon a change in direction. The roughly 50 µm amplitude oscillatory behavior observed during motion correlates with the 1mm pitch of the lead screw. The average repeatability for each axis was about ±5 µm (Figure 2C). Real velocities aligned closely with commanded velocities (Figure 2D). For the remainder of our experiments in this manuscript, we have not compensated for this thread backlash.

**Figure 2:**
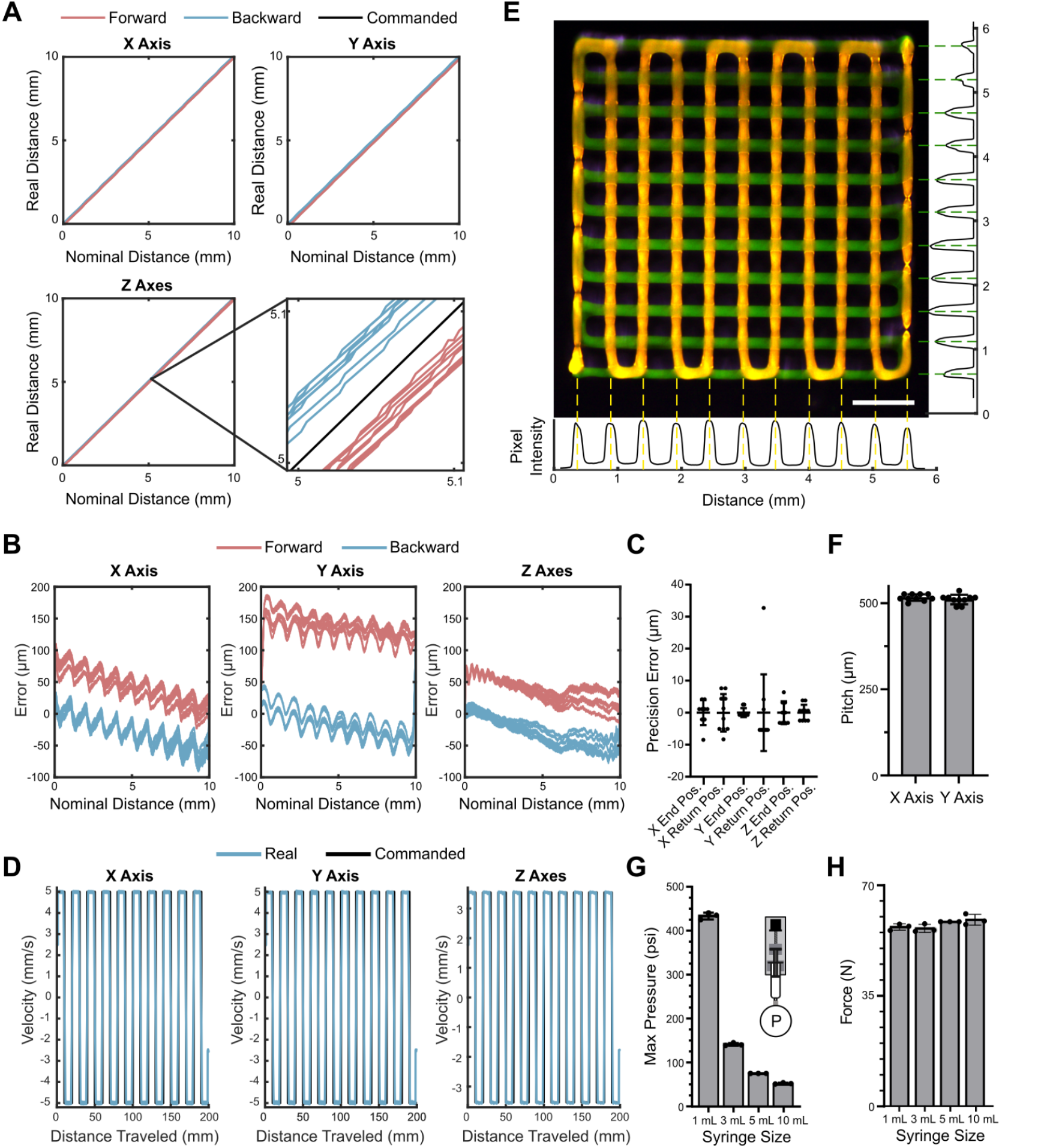
Linear Actuator Characterization and Multimaterial Printing. Precision and accuracy of the x-, y-, and dual carriage z-axes were analyzed by commanding a 10 mm forward-and-backward sawtooth motion for 10 cycles and continuously measuring real position using an external optical linear encoder. **A)** Nominal distance plotted against real distance over 10 cycles. **B)** Absolute error of real distance from nominal distance plotted along the nominal path over 10 cycles. **C)** Encoder-measured positions at the end and after returning from each cycle for each axis. Each group of locations was centered by their mean to assess precision. Average standard deviation, 4.9 µm. **D)** Real velocities were calculated from the derivative of the encoder data smoothed over a window of 500ms and plotted against commanded velocities. **E)** Multimaterial crosshatch printed with green and red fluorescent pluronic inks. Average pixel row and column intensity is plotted across the x-axis (red) and y-axis (green), respectively. Scale bar, 1 mm. **F)** Pitch, or distance between filaments, calculated from pixel intensity peaks for x-axis (516 ± 9.50 µm) and y-axis (510 ± 13 µm). Mean ± s.d. **G)** Maximum extrusion pressures achieved in 1 mL (433 ± 8 psi), 3 mL (141 ± 3 psi), 5 mL (75 ± 0 psi), and 10 mL (52 ± 2 psi) syringes. **H)** Maximum forces generated for 1 mL (57 ± 1 N), 3 mL (56 ± 1 N), 5 mL (58 ± 0 N), and 10 mL (59 ± 2 N) syringes. For each syringe size, N=3; mean ± s.d.

We assessed effective printing accuracy by printing a simple two layered crosshatch geometry onto a glass slide using Pluronic F-127 ink with red or yellow-green fluorescent dye (Figure 2E, Video S1). The centroid location of each printed filament along the x- and y-axes was found from analysis in ImageJ, and the average calculated pitch was 513 ± 11.4 µm from a commanded pitch of 500 µm (Figure 2F).

To measure the maximum pressures that the extrusion actuators can generate for printing viscous bioinks, luer-lock syringes (BD) were connected to a pressure gauge (Ashcroft, 25W1005) and plunged until the actuators stalled. Maximum pressures for 1 mL, 3 mL, 5 mL, and 10 mL syringes were 433 psi, 141 psi, 75 psi, and 52 psi, respectively, which all correspond to a total force generation of approximately 57.5 N (Figure 2G-H).

### 2.3. Multimaterial Active Mixing Printing

Active mixing nozzles employ a rotating impeller to mix two or more incoming streams of material, and are especially well suited to efficiently mixing thick, non-Newtonian and yield stress fluids which are hard to efficiently mix via passive means.^[19]^ In addition, active mixers can be programmatically turned on and off to create well mixed or two-sided (Janus) filaments. However, these active mixers have previously been applied solely to non-living inks.^[19,35,36]^ Here, we use the Printess to demonstrate an active mixing nozzle capable of real-time active mixing of cell-laden bioinks (**Figure 3A**). The mixing nozzle is mounted onto the Printess and features a 3D-printed printhead and a small rotary motor, powered and controlled from the RUMBA control board on the Printess, to drive a 3D printed impeller that churns two incoming bioink streams together (Figure 3B). The revolutions per minute (RPM) of the impeller and mix ratio between the two inks can be easily and arbitrarily modulated during printing using G-Code commands (Figure 3C). We characterized the ink switching dynamics by printing a snake pattern switching between uncolored and green Pluronic inks in a step and ramp regime (Figure 3D). In the step regime, we observed a 120 µL delay from command to actual material switching, which we subsequently compensated for with an initial purge to demonstrate precise patterning control. Similarly, we observed a 200 µL delay in the ramped regime, which can also be compensated for with an initial purge. We next characterized the nozzle’s mixing efficiency by printing filaments with two Pluronic inks with different fluorescent beads (Figure 3E). Without active mixing, the two inks remain separated as a Janus filament after extrusion, while partial and full mixing can be observed at mixing speeds of 100 RPM and 200 RPM, respectively.

**Figure 3:**
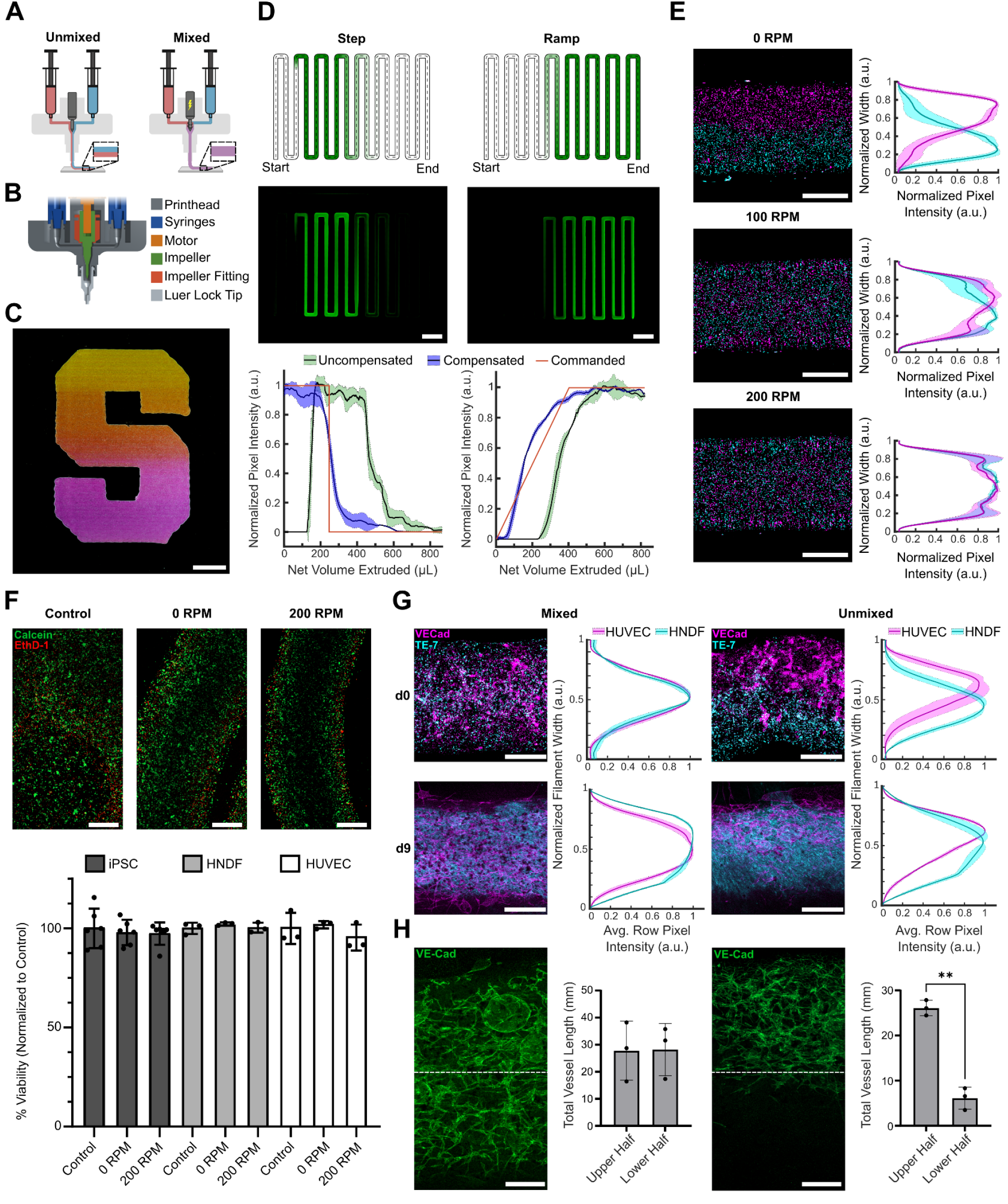
Multimaterial Active Mixing Printing. **A)** Schematic of two-material filament printed with the motor off (left) and motor on (right). **B)** Computer-aided design (CAD) rendering of mixing nozzle printhead with its main components. **C)** Stanford “S” logo printed as gradient between two Pluronic F-127 inks with different fluorescent dyes. Scale bar, 1 cm. **D)** Snake patterns printed using colorless and green fluorescent Pluronic F-127 inks. Nozzle was primed with the colorless ink, and the green ink was printed via a commanded step (left) or ramp (right). After analysis of uncompensated printing dynamics, print was repeated with purge compensation to reduce the effect of the nozzle dead volume. Schematic (top) and photographs (middle) of uncompensated print. Graph of the commanded material-switching profile and the pixel intensities along the printed snake profiles as shown by the dotted line in the schematic (bottom). Scale bars, 1 cm. **E)** Confocal images of filaments printed using Pluronic F-127 with two different fluorescent beads (left) and corresponding average row pixel intensities (right). Mean ± s.d. is plotted. N=5. Scale bars, 1 mm. **F)** Confocal images of induced pluripotent stem cells (iPSCs) printed with the motor on (200 RPM) and off (0 RPM) and casted as control with viability stain (top). Viability of printed iPSCs, human neonatal dermal fibroblasts (HNDFs), and human umbilical vein endothelial cells (HUVECs). Scale bars, 500 µm. **G)** Confocal images of coprinted and cultured HUVECs and HNDFs, both with (left) and without (right) active mixing, stained at day 0 and day 9 of culture with vascular endothelial cadherin (VE-Cad) primary antibody and Fibroblasts Antibody (TE-7), respectively. Graphs of the average row pixel intensities of each stain across the width of the printed filament convey cell distribution. Mean ± s.d. is plotted. N=3. Scale bars, 500 µm. **H)** Confocal images of HUVECs stained with VE-Cad after day 9 of culture following mixed (left) or unmixed (right) printing with HNDFs. Analysis of total vessel length of microvascular networks in the upper versus lower halves of the filament show no significant difference in the mixed condition and a significant difference (p=0.0032) in the unmixed condition via paired t-test. N=3. Scale bars, 250 µm.

We then demonstrated that the active mixing nozzle could be used to print human induced pluripotent stem cells (hiPSCs), human dermal fibroblasts (HNDF), and human umbilical vein endothelial cells (HUVEC) using gelatin-fibrinogen bioinks. The mixed condition (200 rpm) yielded printed cells exhibiting high viability compared to casted (non-printed) control (>95%) (Figure 3F). To assess the effect of active mixing on printed and cultured tissues, we co-printed and cultured HUVECs and HNDFs in both mixed and unmixed conditions, staining immediately post-printing and after 9 days with fibroblast marker TE-7 and endothelial specific adhesion molecule vascular endothelial (VE)-cadherin (Figure 3G). To track cell migration across the mixed or Janus filaments, the average row pixel intensities of the fluorescent channels were assessed along the filament width. In both the mixed and unmixed conditions, the fibroblasts had spread across the full width of the filament, supporting highly migratory behavior.^[37]^ In contrast, the endothelial cells had undergone vasculogenesis throughout the width of the mixed filament, but the Janus (unmixed) vascular networks were found only towards the side on which endothelial cells were initially printed (Figure 3G-H and S3). Prior studies have demonstrated robust endothelial cell angiogenesis towards a spatially separated source of fibroblasts.^[38,39]^ Given the rate at which fibroblasts can migrate through fibrin filament, the spatial separation of HUVECs and HNDFs in our printed filament and potential biochemical gradients is quickly lost, and the densely compacted fibroblasts may be impeding efficient angiogenic sprouting. Overall, these data confirm that active mixing nozzles can effectively mix living bioinks at a shear rate compatible with high cell viability for multimaterial tissue printing.

### 2.4. Multimaterial Multinozzle Printing

Multimaterial multinozzle 3D (MM3D) printheads enable high throughput and versatile printing, with broad applications in robotics, origami metamaterials, and bioprinting. However, previous MM3D printheads utilized precise pressure control and a custom motion control stage, costing over $300,000.^[9]^ To facilitate broader adoption of this method, we have developed and characterized an MM3D printhead module that is compatible with the Printess platform, and editable MM3D printhead files have been made available online. The MM3D printhead is printed via SLA and features two parallel channel networks that ultimately combine at an array of eight nozzles, greatly increasing throughput of multimaterial printing (**Figure 4A**). This nozzle is mounted onto the Printess using a dual z-axis multinozzle backmount. Flow consistency across each of the eight nozzles is critical and dictated by the uniformity of flow path resistance and, ultimately, the SLA print-fidelity of the MM3D printhead. This consistency can be tuned to accommodate different SLA printers by modifying the diameters and lengths of each nozzle individually. The multinozzle used in these experiments has a flow deviation of less than 10% across all nozzles as measured by extruded mass of Pluronic (Figure 4B). A powerful feature of MM3D printheads is their ability to rapidly switch extruded materials within a single filament. Using the low cost syringe pump linear actuators of the Printess, we tested various material switching frequencies, demonstrating a consistent resolution down to 1 mm (Figure 4C-D), which, at a print speed of 5 mm/s corresponds to a 5 Hz switching frequency. While this switching frequency is below the 50 Hz achieved previously in MM3D using rapidly switching pressure controllers,^[9]^ the results here are generated with a homemade $30 syringe pump.

**Figure 4:**
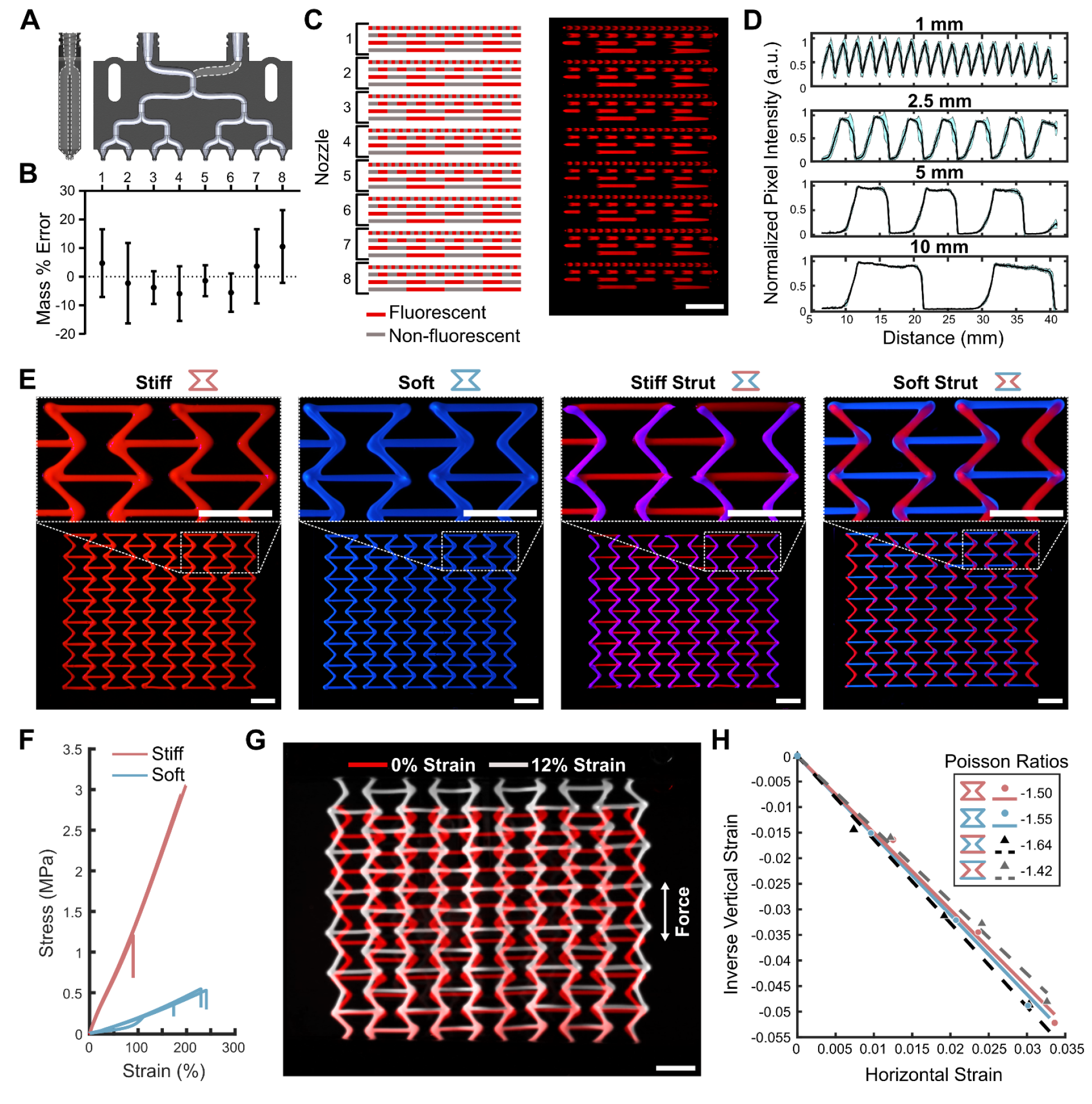
Multimaterial Multinozzle Printing. **A)** Side view (left) and front cross-section view (right) of multinozzle CAD rendering. **B)** Analysis of flow uniformity across each of the eight outputs as measured by mass of Pluronic F-127 extruded. N=4. **C)** Schematic (left) and photograph (right) of a four-filament array alternating between colorless and red Pluronic inks at switching distances of (in descending order) 1 mm, 2.5 mm, 5 mm, and 10 mm. Scale bar, 1 cm. **D)** Analysis of pixel intensity across each filament row. Mean ± s.d. N=8. **E)** Photographs of re-entrant honeycomb auxetic structures printed from stiff and soft silicones. Printing time for each structure was 2 minutes. Scale bars, 1 cm. **F)** Stress-strain curves for stiff (1.247 ± 0.034 MPa) and soft (0.135 ± 0.011 MPa) silicones from tensile testing. Mean ± s.d. N=3. **G)** Overlaid photographs of unstrained and strained stiff auxetic structures. Scale bar, 1 cm. **H)** Horizontal versus vertical strains of each auxetic structure during tensile testing and corresponding Poisson ratios.

One application of this multinozzle is the printing of multimaterial auxetic structures, whose frequently periodic structure is amenable to parallelized manufacturing with MM3D printheads. Auxetic lattices have diverse functions, including as biomimetic support devices, wearable electronics, and mechanical grippers.^[40–43]^ While these are typically manufactured using a single material, multimaterial auxetic lattices manufactured with a combination of stiff and soft struts can exhibit improved auxetic performance.^[43–47]^ We designed a re-entrant honeycomb auxetic structure and printed four versions with the horizontal struts and the oblique legs composed of different combinations of stiff and soft silicones (Figure 4E, S4). Uniaxial tensile testing showed about an order of magnitude difference in Young’s modulus between the stiff (1.247 ± 0.034 MPa) and soft (0.135 ± 0.011 MPa) silicones. We measured the Poisson ratio of each auxetic structure from video analysis during uniaxial tensile testing and demonstrated that the Poisson ratio can be tailored by material selection alone with no change in geometry (Figure 4G, 4H, Video S2). A lower ratio of stiffness between the horizontal struts and oblique legs corresponded to a lower Poisson ratio. Overall, the Printess MM3D platform printed these auxetic lattices in a single parallelized pass in approximately 2 minutes, supporting higher-throughput and multimaterial printing methods.

### 2.5. Embedded and Multimodal Printing

Embedded 3D printing into a viscoplastic and self-healing support bath can allow for freeform 3D writing of soft materials.^**[5**,**48**,**49]**^ To demonstrate and characterize embedded printing on the Printess, we printed a variety of objects at different scales. At large scale, we printed a 7 cm-long dragon using Carbopol with pigment (**Figure 5A**). To further quantify accuracy, we then printed a 2 cm-long Stanford bunny using carbopol with lithium bromide as a contrast agent (Figure 5B, Video S3). We took micro-computed tomography (Micro-CT) scans of the printed bunny to compare with the model used for G-code generation and calculated a Hausdorff Distance of 0.25 ± 0.34 mm (mean ± RMSE) (Figure 5C). Most of the deviation from the model occurred when the extruders made frequent jumps and retractions between the ears due to layer-by-layer slicing. This error could be reduced with the use of alternative, non-layer-by-layer slicing algorithms that minimize these jumps.

**Figure 5:**
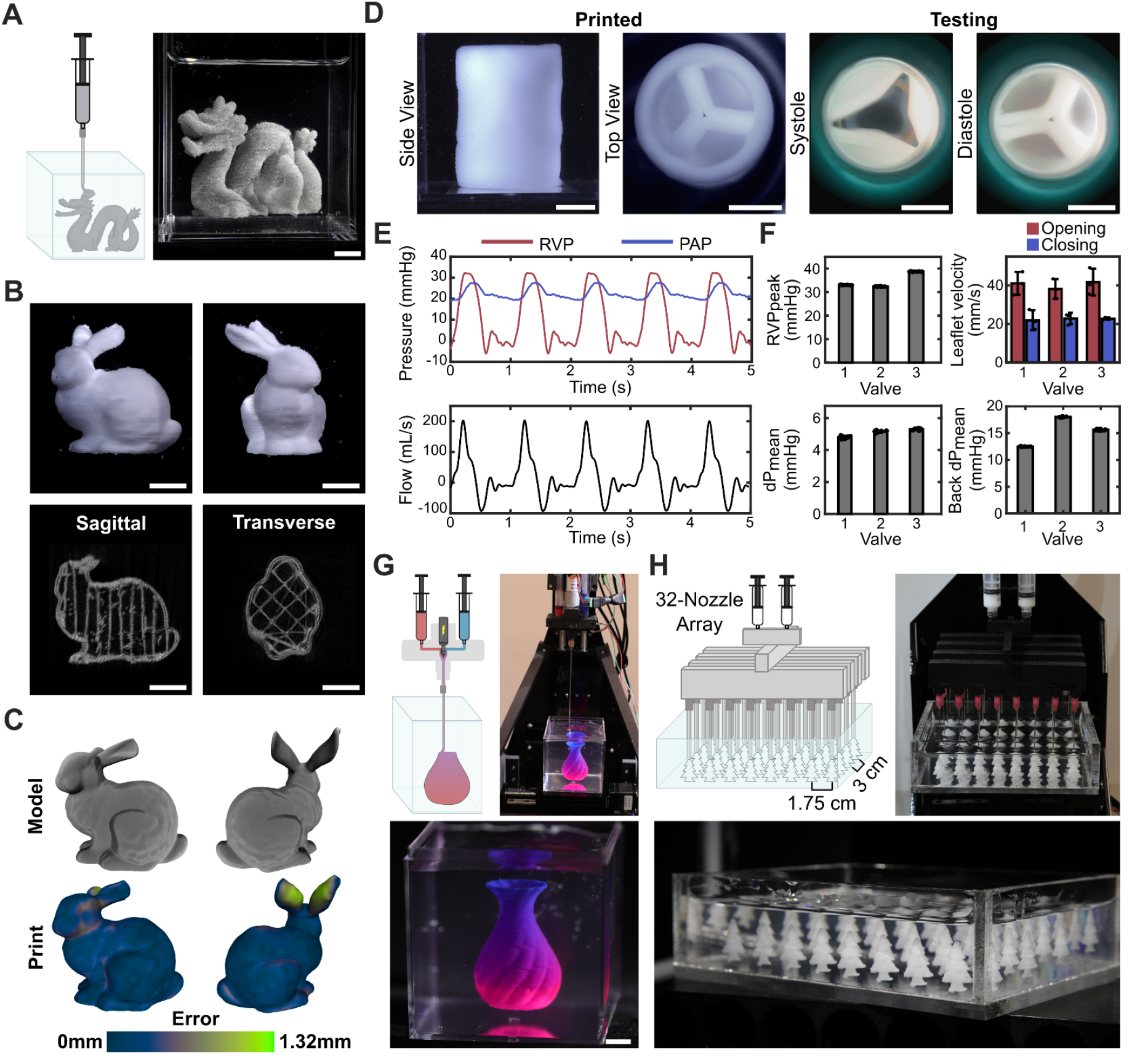
Embedded and Multimodal Printing. **A)** Schematic and photograph of embedded print of a dragon using carbopol mixed with pigment. Scale bar, 1 cm. **B)** Photographs (top) and Micro-CT scans (bottom) of the Stanford bunny printed with infill using carbopol mixed with lithium bromide. Scale bar, 5 mm. **C)** Model (top) and reconstructed Micro-CT scan (bottom) of the Stanford bunny with measured deviation (0.25 ± 0.34 mm). Hausdorff Distance mean ± RMSE. **D)** Photographs of a trileaflet valve and root after printing using PEGDA and carbopol (left) and while in systole and diastole during *in-vitro* hemodynamic testing in a univentricular simulator (right). Scale bar, 1 cm. **E)** Representative continuous pressure and flow data from valves during hemodynamic testing. RVP = Right Ventricle Pressure; PAP = Pulmonary Artery Pressure. **F)** Peak right ventricle pressures, mean transvalvular pressures (dP), mean back-pressures (Back dP), and opening and closing leaflet velocities for three valves. N=8-10 cycles for peak and mean calculations. N=3 leaflets for velocities. Mean ± s.d. **G)** Schematic and photographs of the multimaterial mixing nozzle embedded printing a vase with gradient between carbopol mixed with red or blue pigment. Scale bar, 1 cm. **H)** Schematic and photographs of a multimaterial 32-multinozzle embedded printing a ‘forest’ of trees made of carbopol mixed with pigment. Print duration was 3.5 minutes. All structures were printed into a 0.1% ETD 2020 Carbopol support bath. For the dragon, vase, and forest, the Printess was modified with higher-mounted extruders by adding mounting holes higher on the acrylic backboard.

To illustrate functional embedded printing, we designed, printed, and hemodynamically tested an idealized trileaflet valve and root of 25 mm inner-diameter. Several examples of fabricated valves using various 3D printing methods and materials have been previously presented,^[50,51]^ and we demonstrate a comparable valve that can be generated on our low-cost platform. We printed our valve using carbopol with PEGDA (Figure 5D). Post-UV curing, the valve was mounted in a univentricular simulator for hemodynamic testing,^[52]^ which demonstrated adequate performance of the 3D-printed valve under native pulmonary valve conditions (Video S4). Figure 5D illustrates co-axial images of the valve in the open and closed states at peak-systole and end-diastole, respectively. Superimposed right ventricular and pulmonary artery representative pressure and pulmonary artery flow waveforms over five consecutive cycles confirm appropriate valve opening with effective forward flow throughout the cardiac cycle (Figure 5E). For an average cardiac output of 2.4 L/min (heart rate = 60 bpm; stroke volume = 46.1 mL) (Figure S5A) and peak-systolic right ventricular pressure of 34.7 mmHg (Figure 5F), the valve effective orifice area was 1.7 ± 0.4 cm^2^ (Figure S5B), and the mean and maximum transvalvular pressure gradients were 5.1 ± 0.3 mmHg and 10.7 ± 1.8 mmHg, respectively (Figure 5F, S5C). Mean back-pressure during diastole was 15.4 ± 2.8 mmHg. Retrograde flow (regurgitant fraction of

28.5 ± 7.7%) (Figure S5D) was likely due to perivalvular leak at the suture points, as appropriate coaptation of the leaflets was observed (Figure 5D). Leaflet motion analysis showed that opening and closing velocities are consistent across leaflets within each printed valve and across valves (Figure 5F and S5E). Further, leaflet closing velocities (22.5 ± 2.9 mm/s) are consistently lower than opening leaflet velocities (40.3 ± 5.8 mm/s), which is in line with physiologic behavior.

Finally, we demonstrated novel multimodal printing by combining embedded printing with multimaterial active mixing and multinozzle printing. We printed a vase with color gradient using our active mixing printhead attached to a long nozzle for embedded printing using carbopol with red or blue pigment (Figure 5G). We also designed and 3D printed a high-throughput 32-multinozzle array, which combines ink flows from two parallel syringes and then splits the flow, using bifurcating trees, four ways in *x* before splitting each flow eight ways in *y*. Each path terminates in a male Luer connector for attaching a long metal nozzle for embedded 3D printing. Using this high-throughput nozzle, we printed a ‘forest’ of 32 tree models in 3.5 minutes (Figure 5H, S6, and Video S5). Using a single nozzle, this print would have required 2 hours. Future applications of these new printing modalities include using multiple cell- and matrix-based biomaterials to generate tissue constructs or support baths with variable compositions and properties as well as for high-throughput generation of small-scale tissue arrays.

## 3. Conclusion

We developed an open-source, low-cost multimaterial 3D printer for use in research laboratories and educational settings. It is built from scratch using stepper motors and off-the-shelf materials, has highly customizable components, including syringe mounts and platforms, and is exceptionally low-cost at approximately $250. We demonstrate its capability of several printing modalities, including multimaterial, multimaterial mixing, multimaterial multinozzle, and embedded. In conjunction with commercially available slicing software, it can print arbitrary 3D models and reconstructed scans. The Printess has also been successfully integrated into the Stanford Bioengineering undergraduate and graduate curriculum, demonstrating its potential in educational settings.

Many opportunities exist for expansion of the bioprinter’s capabilities and applications. We plan to continue development of these printing modalities to study gradients in cell type and matrix environment towards printing and maturing functional tissues. The Printess contributes to broadening the accessibility of bioprinting with the long term aim of furthering the field of tissue engineering. Lowering the cost of bioprinters is an important step toward achieving broad adoption of biofabrication techniques across disciplines.

## 4. Experimental Section

### 4.1. Printess Materials and Methods

All CAD files, firmware, and scripts are available open-source on Zenodo at https://doi.org/10.5281/zenodo.13173619. The latest versions of these files and the construction and operation manual (Supporting Methods) can be found on Github at: https://github.com/weiss-jonathan/Printess-Low-Cost-3D-Printer.

#### 4.1.1. 3D Printed and Laser Cut Components

CAD files were developed in SolidWorks or OnShape. All experiments in this work were conducted on a Printess that was constructed using components printed on a Mark II 3D printer (MarkForged, Watertown, MA) using black onyx and carbon fiber filament. The Printess has also been constructed using parts printed using PLA on a Prusa I3 MK3S 3D printer. Black acrylic sheets (12’’ x 12’’ x 0.25’’) were laser cut using a Glowforge laser cutter to create the base and back plates. The mixing nozzle and multinozzle printheads were printed using Biomedical Black resin on a Formlabs 3B printer.

#### 4.1.2. Stepper Motors

All motion on this printer is driven by six NEMA 11 linear stepper motors. Two 100 mm motors move the printing stage in the X and Y directions, two 100 mm motors independently raise and lower each extruder in the Z direction, and two 50 mm motors independently control extrusion from two syringes. 3D printed components and M3 screws secure each motor to the back and base acrylic plates. Precision and accuracy of motors was measured using an optical linear encoder (US Digital; EM2-0-2000-N) with a 2000 lines per inch strip (US Digital; LIN-2000-1-N). The strip was mounted directly to the stepper motor table, the linear encoder was mounted alongside the table, and a 10 mm sawtooth motion with 10 cycles was commanded. Data was collected using Automation1 (Aerotech) software.

#### 4.1.3. Syringe Mounts

Syringe mounts were designed to accommodate 1 mL, 3 mL, 5 mL, and 10 mL BD plastic syringes. Independent control of two mountable syringes with interchangeable nozzles enables simultaneous multimaterial printing of soft materials. To prevent finger injuries from the stepper motors if the printers are used for educational purposes, we provide designs of 3D-printable protective cuffs that prevent the linear stepper motor tables from reaching either end of the rail.

#### 4.1.4. Control Board

The RUMBA+ board, a RepRap board with an onboard ATmega2560, can host 6 stepper motors and drivers. TMC2208 drivers were used for the experiments in this manuscript. For safer printer operation, the stepper motor drivers should be tuned using the on-board trimpots to limit the force of the linear stages to a level just sufficient for operation. The RUMBA+ board possesses a number of features including multiple fan pins, temperature connectors, a port to use with touchscreen controllers, and end stop connections, all of which may be used in customization of the bioprinter. The RUMBA+ uses a mini USB connection, and wiring of the stepper motors is described in the Supporting Methods.

#### 4.1.5. Firmware and Software

The open-source 3D printer firmware Marlin was used with the RUMBA+ board. It was modified to designate the extrusion axes as additional linear axes that can be controlled independently from the other standard X,Y, and Z linear axes (Supporting Methods). A mac or windows laptop was used to interface with the board using the open-source graphical user-interface software Printrun (Pronterface). The modified version of Marlin is available in the Zotero repository. 3D test models, including the Stanford dragon, Stanford bunny, and vase, were sliced into G-code using UltiMaker Cura v5.8.0 slicing software. We provide configuration instructions to set default print parameters within Cura (Supplemental Methods).

### 4.2. Cell Culture

All cells were maintained at 37 °C and 5% CO_2_ in T25 or T75 flasks (Thermo). HNDFs (Lonza; CC-2509) were cultured in DMEM/F12 (Thermo; 11320033) supplemented with 10% FBS (Sigma; F4135) and 1% PenStrep (Fisher; 15-070-063) with media changes every 3 days and passages at a 1:5 ratio when confluent. HUVECs (Lonza; C2519A) were cultured in Endothelial Cell Growth Medium-2 (EGM-2) (Lonza; CC-3162) supplemented with 1% PenStrep with media changes every 2 days and passages at 1:3 ratio when confluent. hiPSCs (SCVI-15; Stanford Cardiovascular Institute Biobank) were cultured in E8 medium with daily media changes and passages every 3 days or when confluent. For passaging, HNDFs and HUVECs were incubated at 37 °C in TrypLE Express (Fisher; 12-605-028) for 3-5 minutes or until fully lifted, washed with equal volume of respective culture media, and centrifuged at 300 x g for 5 minutes before resuspension and seeding. Similarly, hiPSCs were incubated in Gentle Cell dissociation reagent [PBS without calcium and magnesium (Corning; 21-049-CV), 0.5 mM EDTA (Sigma; EDS-500G), 1.8 g/L sodium chloride (Sigma; S7653)] for 8-10 min at 37 °C, resuspended in equal volume Essential 8 (E8) medium (Thermo; A1517001), centrifuged for 300 x g for 5 min, and seeded at 20,000 cells/cm^2^ with 10 µm Y-27632 dihydrochloride (BioGems; 21293823) onto 12 µg/cm^2^ GelTrex (hESC-qualified) (Thermo; A1413302). All cells were tested negative for mycoplasma by PCR every month. Stanford University Cardiovascular Institute Biobank operates under IRB approval and in accordance with the Stanford Stem Cell Research Oversight (SCRO) committee.

### 4.3. Printing, Tissue Culture, and Staining

During printing, nozzle translation speeds were 5 mm/s, and a straight metal nozzle (Nordson) with inner diameter of 0.1 mm (crosshatch; Figure 2E), 0.2 mm (Stanford bunny; Figure 5B), 0.41 mm (gradient vase; Figure 5G), 0.58 mm (cell prints and Stanford dragon; Figure 3F-H and Figure 5A), 0.84 mm (Stanford “S” and snake; Figure 3C-D), 1.36 mm (trileaflet valve; Figure 5D), or 1.6 mm (fluorescent bead; Figure 3E) was used. Cells were printed directly into an untreated 12-well plate (Costar) at cell concentrations of 30 million HUVECs / mL and 10 million HNDFs / mL. Viability controls were extruded manually directly from the syringe without a connected nozzle. After printing, tissues were immediately cast with a 1:1 ratio of HUVEC and HNDF media containing 10 U/mL bovine thrombin (Rocky Mountain Biologicals) and antibiotic-antimycotic (Thermo). Tissues were then incubated at room temperature for 20 minutes to allow for fibrin polymerization. For viability assessment, media was exchanged for a 1:1 ratio of HUVEC and HNDF media with Live/Dead stain (Thermo) at concentrations of 2 µL ethidium homodimer-1 and 0.5 µL calcein-AM per mL of incubation media and incubated at 37°C for 15 minutes prior to confocal imaging. Otherwise for long-term culture, tissues were transferred to a 37 °C incubator for 15 minutes before media was exchanged with a 1:1 ratio of HUVEC and HNDF media with antibiotic-antimycotic and 11 µg/mL bovine lung aprotinin (Sigma). A ¾ media change without aprotinin was performed every 2 days. Tissues were fixed for 20 minutes using 4% paraformaldehyde, permeabilized for 15 minutes using 0.1% (v/v) triton X-100, blocked for 1 hour in animal free blocking solution (Vector Labs), stained overnight using primary antibodies 1:200 VE-Cadherin (Cell Signaling; D87F2; Lot: 6) or 1:200 Fibroblasts Antibody TE-7 (Novus Biologicals; NBP2-50082; Lot: 3990219), and stained overnight again using Alexa Fluor secondary antibodies (Thermo; A21428, Lot: 2272588; A21235, Lot: 2284596).

### 4.4. Ink Preparation

Pluronic inks were made with 35 wt% Pluronic F-127 (Sigma Aldrich) dissolved in DI water at 4°C overnight. Fluorescent inks consisted of 1.5 v/v% either red or yellow-green fluorescent dyes (Risk Reactor) and were mixed using a pipette with the pluronic ink while on ice. Inks were left at room temperature for a few minutes until solidified prior to printing. For the mixing nozzle fluorescent bead print, 400 μL of 10 μm red or yellow-green polystyrene microbeads (FluoSpheres; ThermoFisher) were added to 2 mL of Pluronic on ice and vortexed before loading into the syringe. Confocal images were recolored to magenta and cyan.

Gelatin solution was prepared by dissolving gelatin type B (Fisher) at 15 w/v% in phosphate-buffered saline (PBS) with calcium and magnesium (Corning) and stirring at 60°C for 6 hours before vacuum sterile filtering. Fibrinogen solution was prepared by dissolving bovine fibrinogen (Rocky Mountain Biologicals) at 100 mg/mL in PBS with calcium and magnesium for approximately 2 hours at 37 °C with gentle agitation on an orbital shaker. After passaging, cell inks were prepared by centrifuging at 300 x g for 5 minutes, aspirating supernatant, resuspending in fibrinogen solution, and finally mixing with gelatin solution and PBS to achieve a net 12% gelatin and 20 mg/mL fibrinogen solution. Bioinks were quickly pipetted into a syringe, air bubbles were removed via inversion, and then syringes were placed on ice for 10 minutes to solidify the bioink prior to printing.

Stiff silicone inks were prepared by combining 8 g DOWSIL SE 1700 with 0.8 g curing agent, 2 g Sylgard 184 with 0.2 g curing agent, and 0.22 g red or blue fluorescent pigment (TechnoGlow) followed by mixing in a FlackTek Speedmixer for 1 minute at 250 RPM. Soft silicone inks were prepared by combining 10 g DOWSIL SE 1700 with 0.4g curing agent and 0.24 g blue fluorescent pigment, mixing in a FlackTek Speedmixer for 1 minute at 250 RPM, adding 0.68 g Sylguard 527 Part A and 0.68 g Sylguard 527 Part B, and finally mixing again in a FlackTek Speedmixer for 1 minute at 250 RPM. Stiffnesses of each silicone were calculated from the initial 10% strain measured via Instron uniaxial tensile testing of dogbones cut from cast silicone sheets using a standard-sized cutting die (ASTM D-412-A-IMP).

All Carbopol gels were generated by adding the desired wt% to a beaker of Milli-Q water, stirring overnight until dissolved at room temperature with an overhead impeller at 200 RPM, and finally increasing pH to 7 using 1 M NaOH while stirring. Embedded print support baths were composed of 0.1 wt% Carbopol ETD 2020 polymer (Lubrizol). The inks were composed of 0.5 wt% Carbopol 2984 polymer (Lubrizol) with either 20 wt% barium sulfate (Stanford bunny), 1 wt% iron oxide and 1 wt% barium sulfate (Stanford dragon), or 2 wt% of either blue or red UV fluorescent powder (TechnoGlow) (gradient vase). Pigments were mixed in a FlackTek SpeedMixer for 1 minute at 250 RPM. The trileaflet valve ink was prepared by mixing 2.3 mL Milli-Q water, 2.5 g 3 wt% Carbopol Ultrez 20 polymer (Lubrizol), 5 mL 40 wt% Poly(ethylene glycol) diacrylate (PEGDA) 20k (Polysciences), 200 µL of 5 wt% Lithium phenyl-2,4,6-trimethylbenzoylphosphinate (LAP; Sigma), and 0.2 g barium sulfate (Sigma) in a FlackTek cup and mixing in a FlackTek SpeedMixer for 1 minute at 250 RPM. PEGDA and LAP were dissolved in Milli-Q water overnight on a roller mixer at 4 °C.

### 4.5. Image Analysis

Crosshatch pixel intensity was measured in ImageJ by plotting the average profiles of each filament separately using the red channel for red pluronic and the green channel with red subtracted for the green pluronic. The filament pitch was then calculated in MATLAB R2024b by defining the position of each filament as the midpoint of its 1/e width.

The mixing nozzle snake pattern pixel intensity was similarly plotted using ImageJ. All plots were normalized to that of a fully green-printed snake to control for imaging setup variations and lighting balance. Step regime data was smoothed with a moving median, and ramp regime was smoothed with a moving mean. Distribution plots for mixing bead and HNDF/HUVEC prints were generated in MATLAB by averaging pixel intensity across each row in each respective channel. Data was then smoothed using a moving mean. Live/dead analysis was conducted using the particle counter in ImageJ. HUVEC microvascular analysis was conducted using AngioTool v. 0.6a. Statistics were calculated in GraphPad Prism v10.1.2.

The multinozzle switching frequency photographs were analyzed in ImageJ by averaging pixel profiles of each respective row.

### 4.6. Multinozzle Flow Test and Auxetic Structures

The multinozzle flow test was conducted by extruding 35 wt% Pluronic F-127 through one channel at a time and collecting the extruded outputs in 8 individually tared PCR tubes. Mass of each tube was measured to calculate net extruded mass from each nozzle, and percent error from the mean (about 100 mg) was calculated. Both channels from two different printheads were tested, and all data was averaged together (n=4).

Auxetic silicone structure uniaxial testing was conducted on a custom-built uniaxial strain device. Both ends of the structure were clamped using flat acrylic grippers, and the structure was strained in increments of 1 mm until about 12% strain. Video was taken from above during the test, and strains were measured in ImageJ. The linear regime of the strain curves (up to about 5% strain) was used to calculate the Poisson ratio.

### 4.7. Valve Design and Printing

The valvular leaflet geometry was generated according to Hamid et al.^[53]^ First, an elliptic paraboloid centered at the valvular root’s edge was generated, and paraboloid points on the outside of the root geometry were omitted. A subsegment of the valvular root was then created by two planes intersecting the centerline of the valvular root and separated by 60 degrees in each direction from the center of the paraboloid. All the leaflet points that crossed the generated planes were modified to be on the respective plane with an offset factor added to prevent the leaflets from overlapping and curing together during printing. Finally, the generated leaflet points were symmetrically patterned in a circular manner around the centerline of the valvular root. After printing, the valve was UV-cured in a FormLabs Cure Station for 5 minutes at room temperature and then transferred from the support bath into PBS to match hemodynamic testing saline conditions for 14-18 hours prior to hemodynamic testing. In this time, the valve reaches a steady-state of swelling that results in a roughly 10% size increase. Printing parameters accounted for this swelling.

### 4.8. Valve testing and Hemodynamic Analysis

The 3D printed valve was mounted into an univentricular ex vivo heart simulator (ViVitro Superpump, ViVitro Labs, British Columbia, Canada).^[52]^ The valve was attached to the simulator via 2-0 Ethibond sutures at the base and a ziptie at the distal end. The univentricular heart simulator is made of a linear piston pump and two compliance chambers that recapitulate physiological systemic or pulmonary pressures and flows using saline. The ViVitest Software (ViVitro Labs, British Columbia, Canada) allows for adjustment of hemodynamic parameters, such as peripheral resistance, stroke volume, flow waveforms, and heart rate. In this study, the hemodynamic parameters were chosen to approximate physiologic right-sided pressures and flows (average cardiac output = 2.6 L/min, RVP_max_ = 38.4 mmHg) to evaluate the short-term biomechanics of the 3D-printed valve.

The univentricular heart simulator incorporates 25-mm flow probes to measure pulmonary flow (Carolina Medical Electronics, East Bend, NC) and pressure sensors to measure pulmonary and ventricular pressure (Utah Medical Products, Inc., Midvale, UT). Before each experiment, all sensors were zeroed and reset by exposing the sensors to atmospheric pressure while the flow sensors were calibrated to create a zero-flux environment. High-speed videography was captured from the distal end of the valve at 1057 frames per second with 1280 × 1024 resolution (Chronos 1.4; Kron Technologies, Burnaby, British Columbia, Canada). Leaflet tip opening and closing velocities were calculated using LoggerPro 3.

Hemodynamic parameters for valve testing were calculated on MATLAB R2024b using code previously developed.^[54]^ Right ventricle pressure and pulmonary artery pressure and flow were obtained directly from the heart simulator. Mean back pressure was calculated by averaging the absolute value of the transvalvular pressure gradient during diastole. Metrics of stroke volume, cardiac output, peak-systolic right ventricular pressure, transvalvular pressure gradient, back pressure gradient, regurgitant fraction, and effective orifice area were averaged over eight to ten consecutive heart cycles.

### 4.9. Micro-CT Scanning

A Bruker SkyScan 1276 CT scanner (source voltage = 85 kV; source current = 200 µA) along with reconstruction software NRecon (Micro Photonics) was used to generate a 3D model and cross sections of the printed Stanford bunny. Hausdorff Distance was calculated in 3D Slicer (www.slicer.org).

## Supporting Information

Supporting Information is available from the author.

**Supporting Video S1: Multimaterial Crosshatch Print Supporting Video**

**S2: Auxetic Uniaxial Tensile Testing Supporting Video**

**S3: Embedded Stanford Bunny Print Supporting Video**

**S4: High-Speed Valve Hemodynamic Testing Supporting Video**

**S5: 32-Multinozzle Tree Print**

## Author Contributions

M.A.S.S, J.W., and A.M.B conceived of and led the design and construction of the Printess as well as wrote the initial manuscript. M.A.S.S., J.W., and L.R. revised the manuscript. J.W., A.M.B, T.T., and F.S. contributed to the supporting information. J.W. conducted printing, characterization, and analysis of experiments. F.S., T.T., and S.S. contributed to the design and construction of the Printess and experiments. F.S. created the valve design. D.R. established the formulation for the valve material. A.S. led valve hemodynamic testing under support from M.M. P.C. and M.S. sutured valves for hemodynamic testing. Y.P. provided pressure measurements for hemodynamic testing. R.N. measured and analyzed valve leaflet velocities. L.R. supported and conducted the analysis of hemodynamic testing. K.P. supported early development of the Printess and its introduction to Hyde Middle School in Cupertino, CA.

## Acknowledgements

We would like to thank Kayla Wolf for the supplemental image of a PLA-printed Printess and Stacey Lee, Seo Woo Song, and Qiuling Wang for their support with the cellular printing. hiPSCs were obtained from Joseph C. Wu, MD, PhD at the Stanford Cardiovascular Institute funded by NIH75N92020D00019. Research reported in this publication was funded by a Chan-Zuckerberg Biohub Investigator Award and by the National Heart, Lung, And Blood Institute of the National Institutes of Health under Award Number DP2HL168563. We would also like to thank the Stanford BIOME Ideathon for funding the original prototype, the Stanford Center for Innovations in In Vivo Imaging (SCi3) small animal imaging center for the Micro-CT scanner, and the Stanford Nano Shared Facilities (SNSF), supported by the National Science Foundation under award ECCS-2026822, for the Instron. J.W. was funded by Stanford Bio-X. J.W., A.M.B., and S.S. were funded by the National Science Foundation Graduate Research Fellowship Program. S.S was funded by the Stanford Graduate Fellowship. L.R. was funded by the Burrough Wellcome Funds Career Awards at the Scientific Interface. Some of the diagrams in Figure 1, 3, and 5 were created in BioRender.com. Stanford dragon and bunny models were from The Stanford 3D Scanning Repository (graphics.stanford.edu/data/3Dscanrep/), and the vase was from GrabCAD user Navaneethan S. (grabcad.com/library/flower-vase-103).

## Data availability statement

All CAD files, firmware, and scripts are available open-source on Zenodo at https://doi.org/10.5281/zenodo.13173619. The latest versions of these files and the construction and operation manual (Supporting Methods) can be found on Github at https://github.com/weiss-jonathan/Printess-Low-Cost-3D-Printer.

