## Supplementary figures and images for "A low-cost, open-source 3D printer for multimaterial and high-throughput direct ink writing of soft and living materials"

### 1 Overview.png

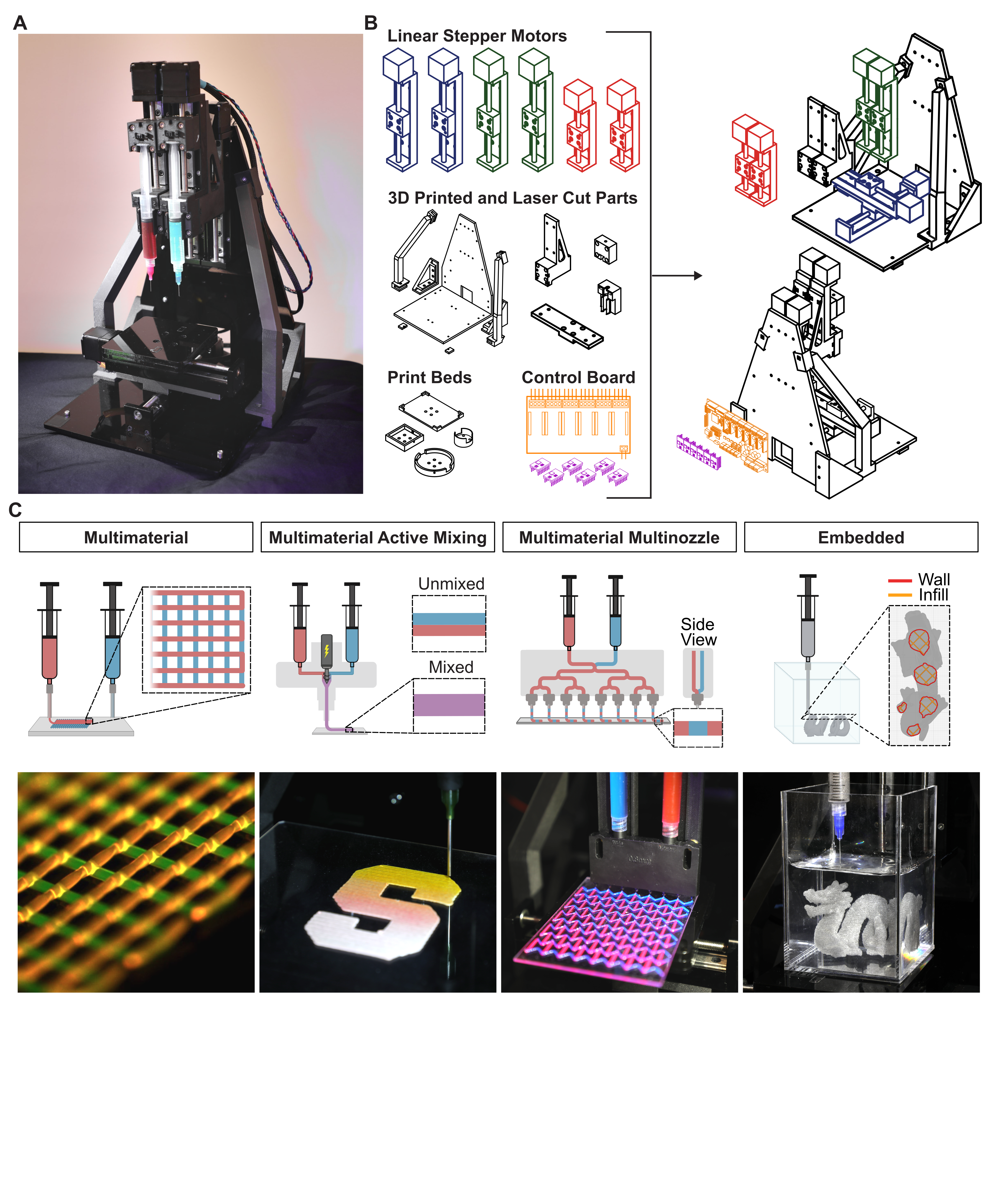

### 2 Characterization.png

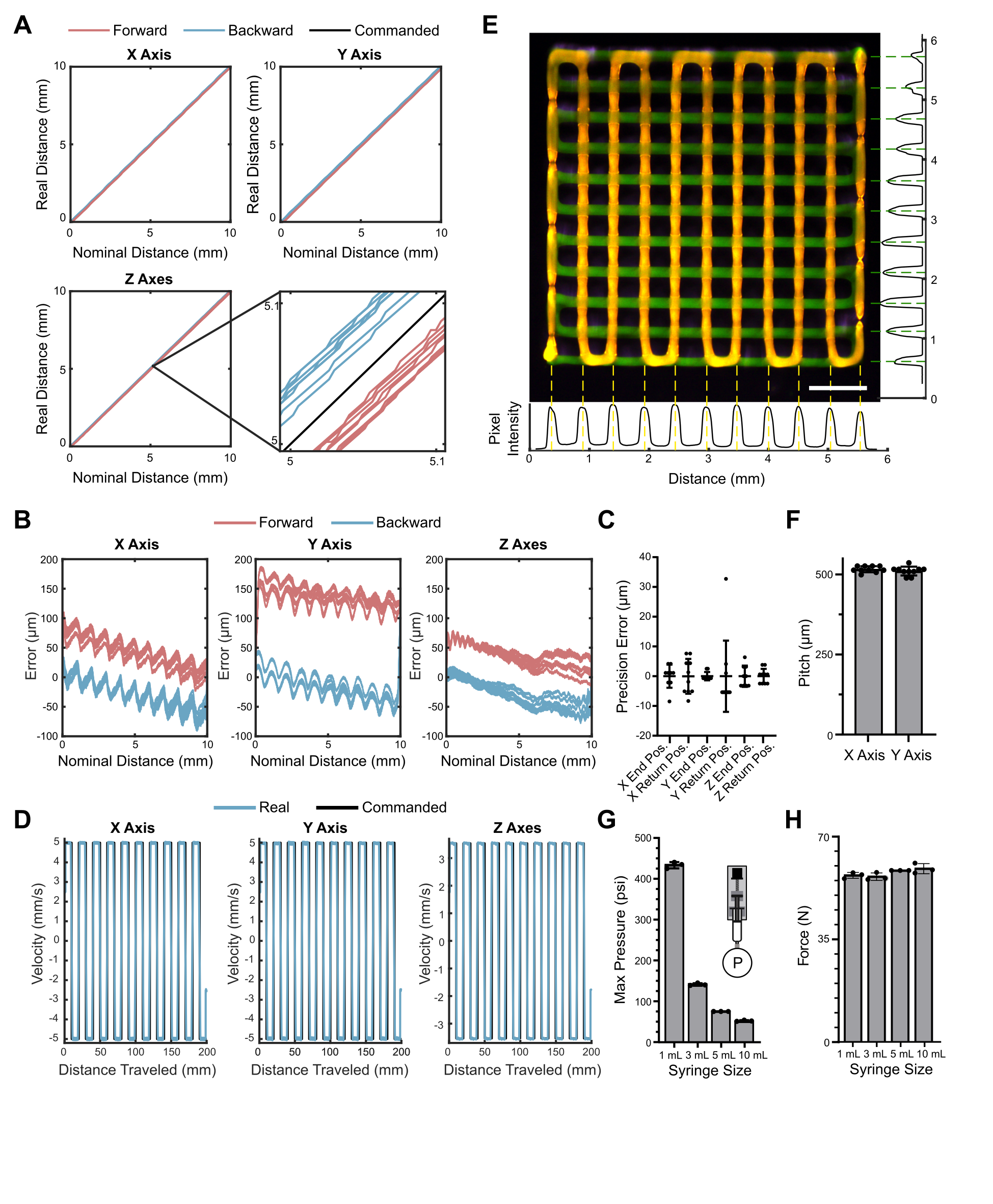

### 3 Mixing.png

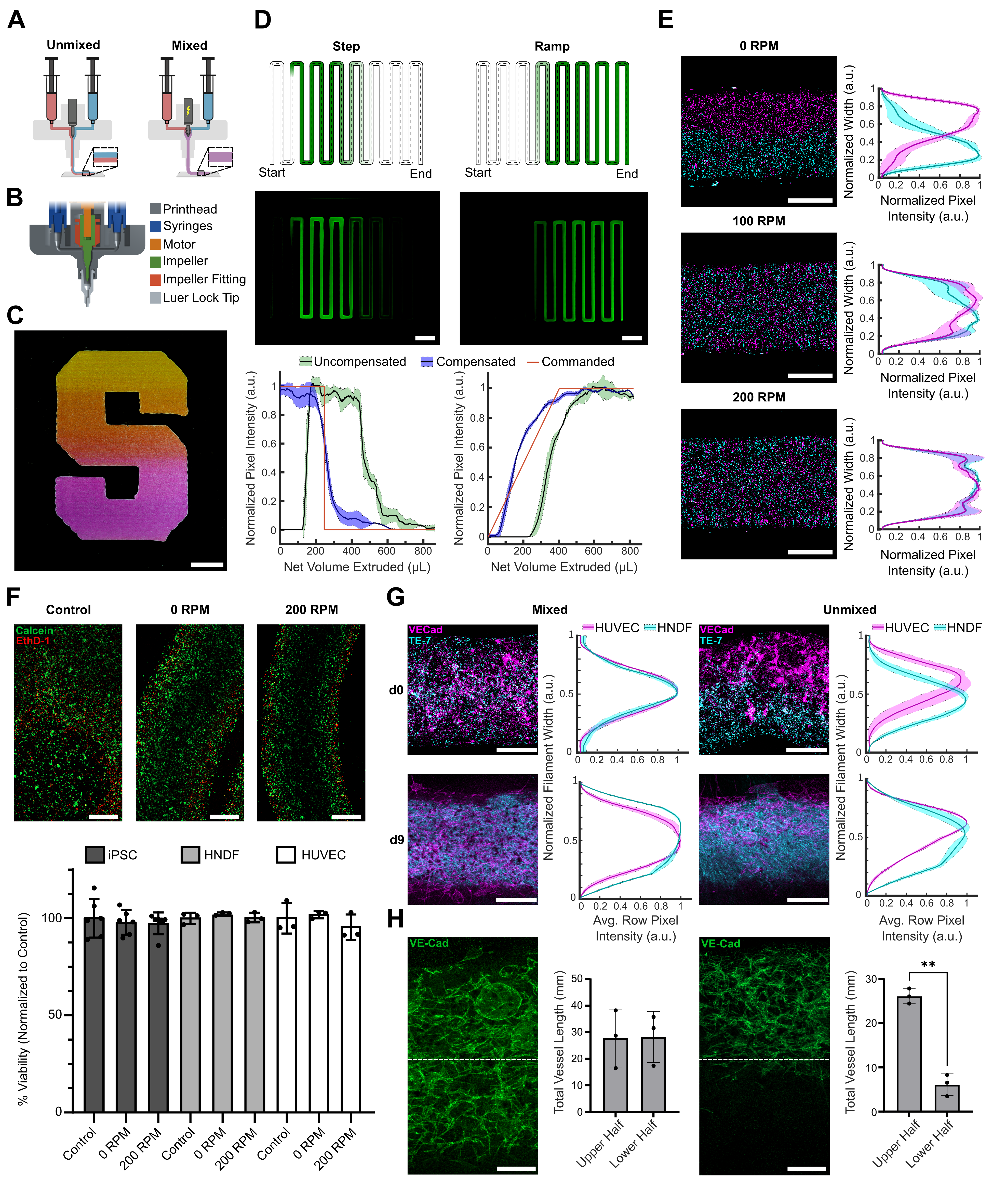

### 4 Multinozzle.png

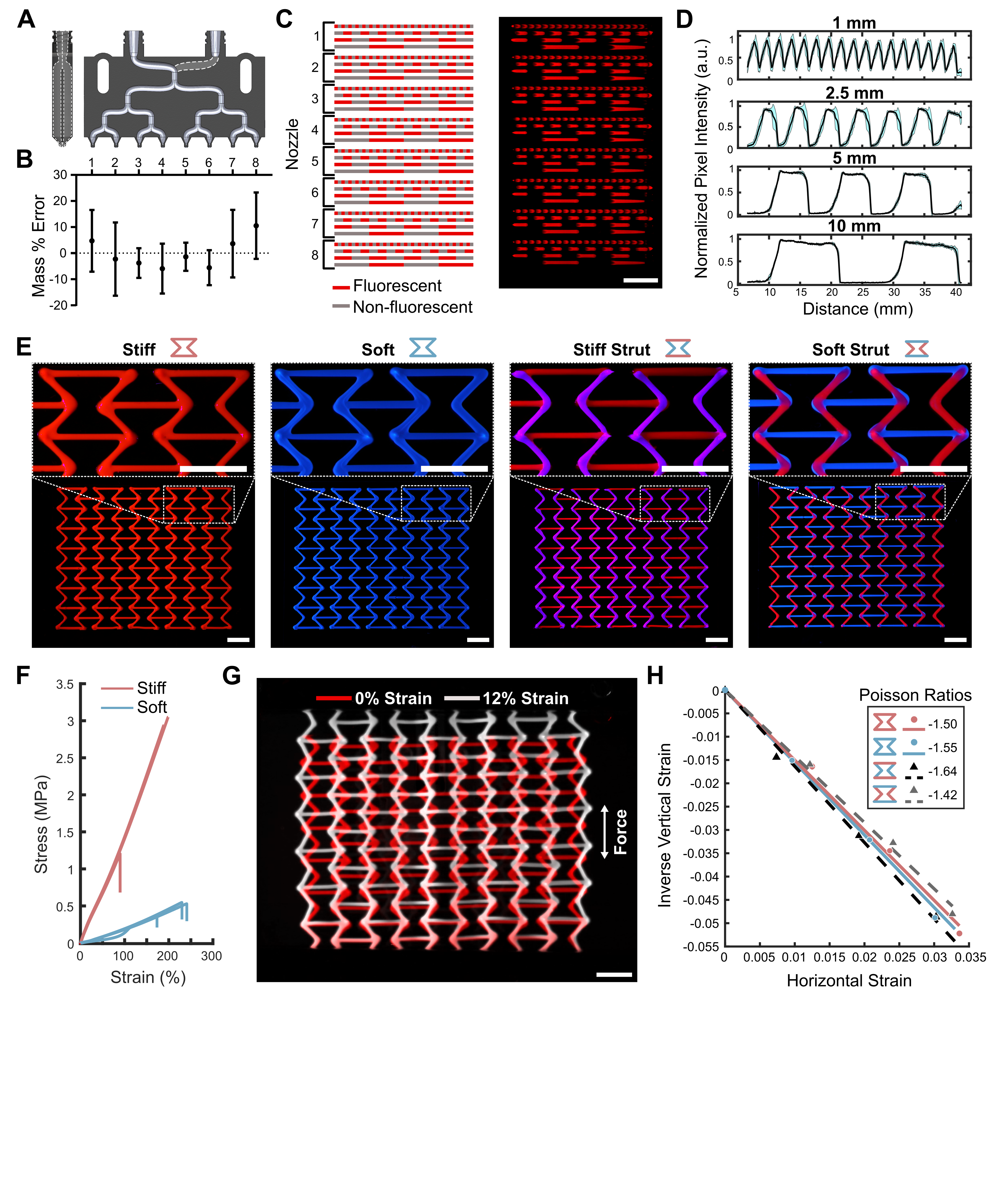

### 5 Embedded Figure.png

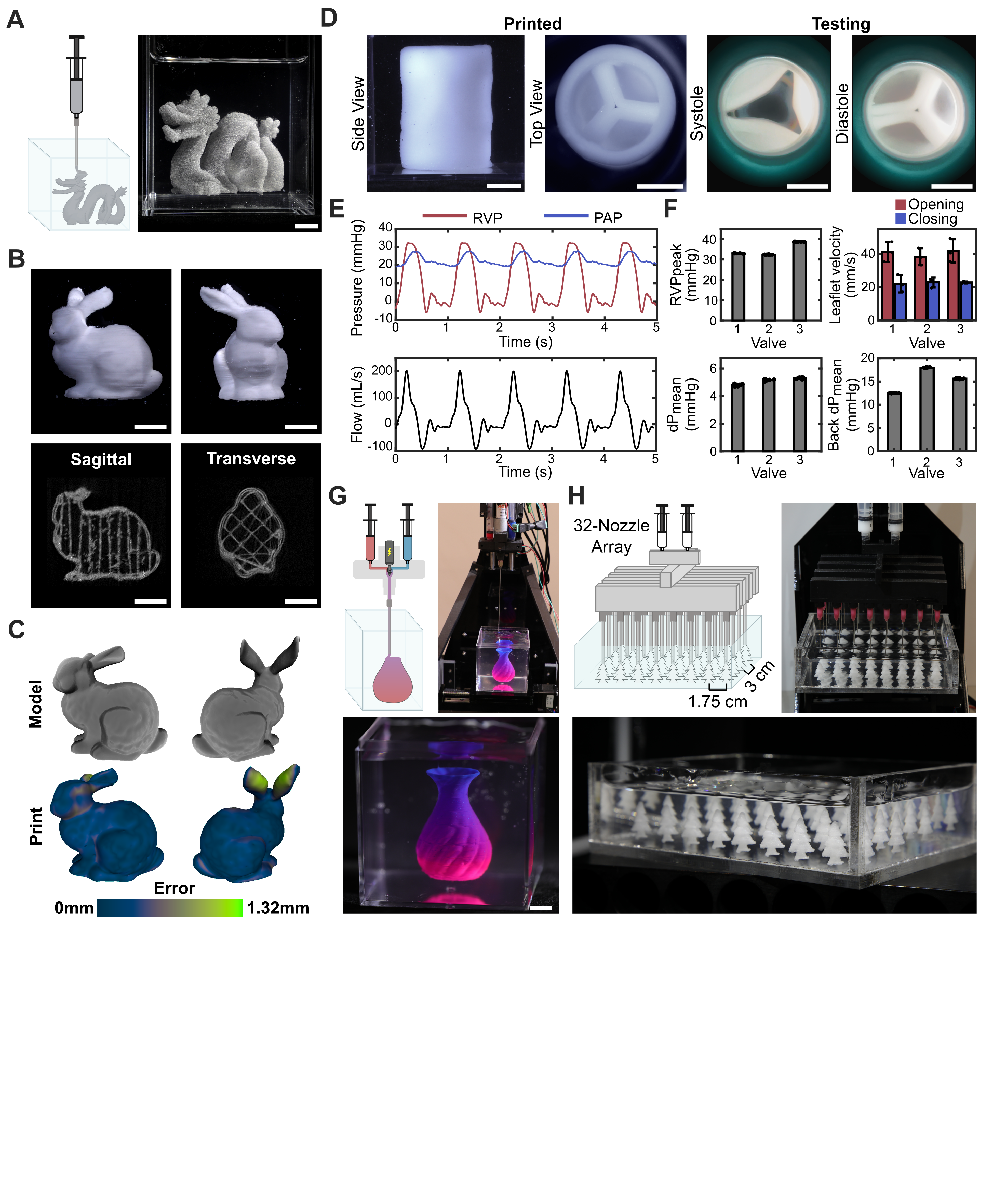
