## Supporting Figures and Methods for "A low-cost, open-source 3D printer for multimaterial and high-throughput direct ink writing of soft and living materials"

**This PDF includes:**

Supporting Figure S1: Printess built in PLA and carbon fiber reinforced-nylon
Supporting Figure S2: Fleet of 6 Printesses
Supporting Figure S3: Endothelial Cell Network Analysis
Supporting Figure S4: Multimaterial multinozzle unit cell
Supporting Figure S5: Trileaflet valve hemodynamic testing
Supporting Figure S6: 32-Multinozzle Printhead
Supporting Table S1: Existing low-cost bioprinters
Supporting Table S2: Bill of materials
Legends for Supporting Movies S1-S5
Supporting Methods: Printess Manual

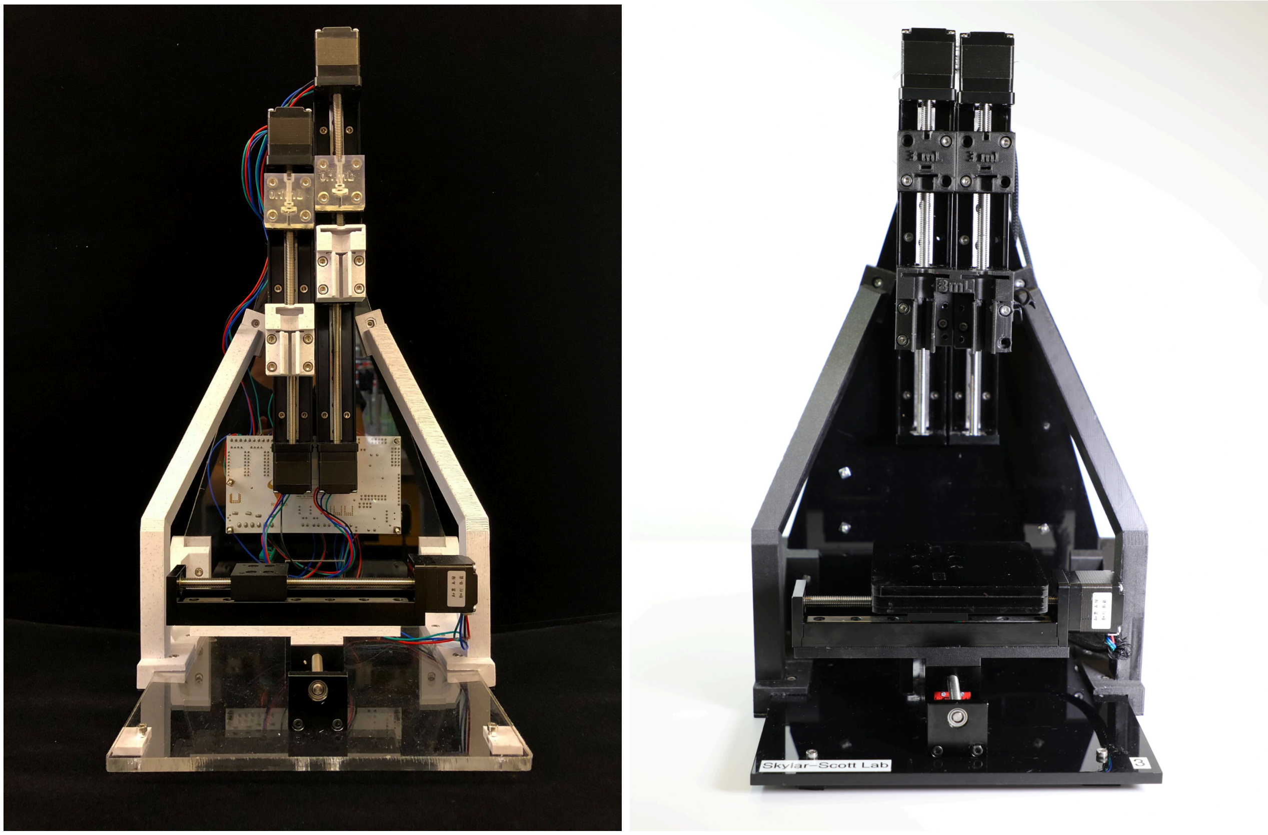

**Figure S1. Printess built using components printed in PLA on a Prusa I3 MK3S (left) and carbon fiber reinforced-nylon on a Markforged Mark 2 (right).**

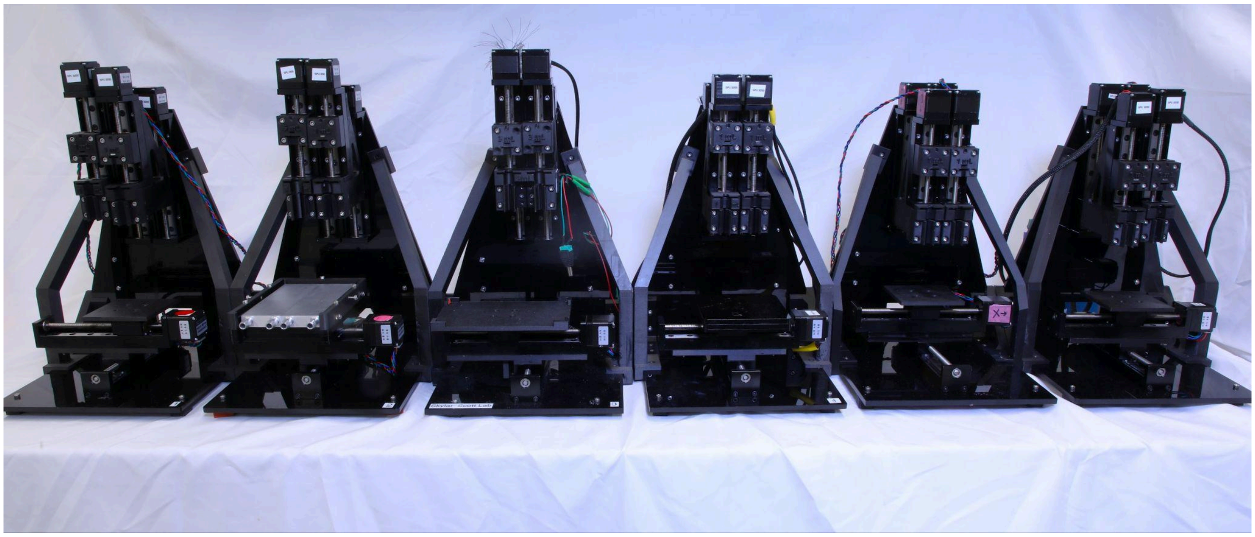

**Figure S2. Fleet of 6 Printesses.** Printers were built using components printed on a Markforged Mark II printer with carbon fiber-reinforced nylon (Onyx). Build-bed types as pictured from left to right: basic flat square platform, custom cooling bed, 6-well plate holder, acrylic basic flat square platform, basic flat square platform, and basic flat square platform.

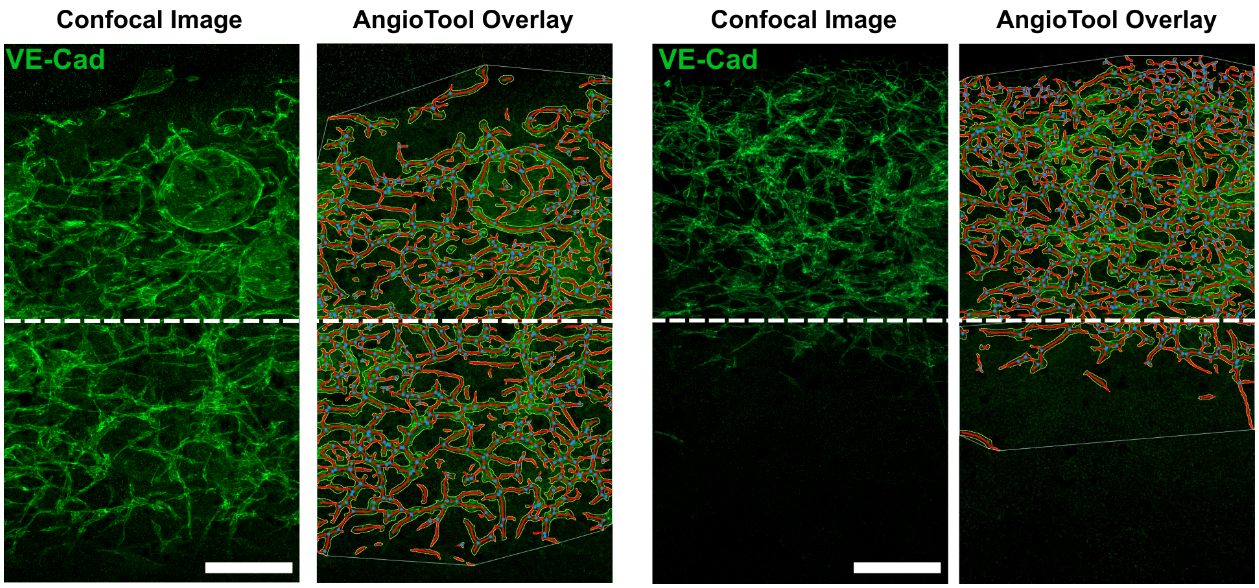

**Figure S3. Endothelial Cell Network Analysis.** AngioTool analysis for calculation of total vessel length in Figure 3H. Scale bars, 250 μm.

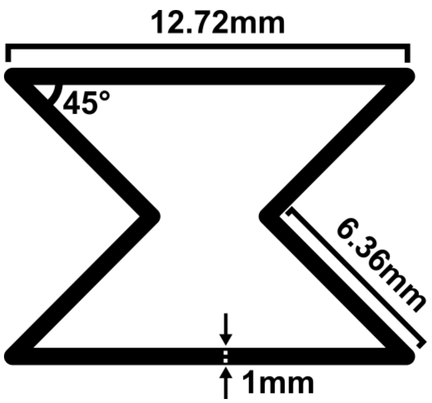

**Figure S4. Dimensions of unit cell for multimaterial multinozzle auxetic structures.**

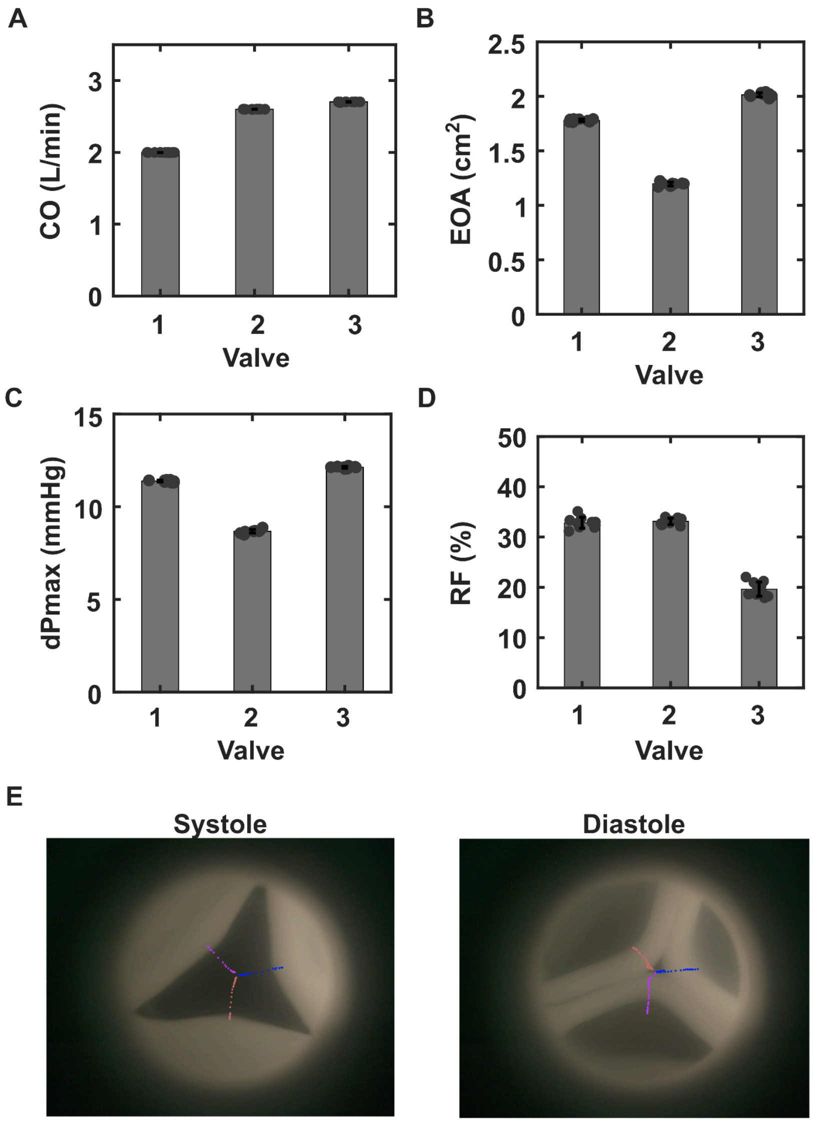

**Figure S5. Trileaflet valve hemodynamic testing. A)** Prescribed cardiac output. N=8-10 cycles. Mean ± s.d. **B)** Effective orifice area. N=8-10 cycles. Mean ± s.d. **C)** Peak transvalvular pressure. N=8-10 cycles. Mean ± s.d. **D)** Regurgitant fraction. N=8-10 cycles. Mean ± s.d. **E)** End-frames of leaflet velocity tracking for valve 1.

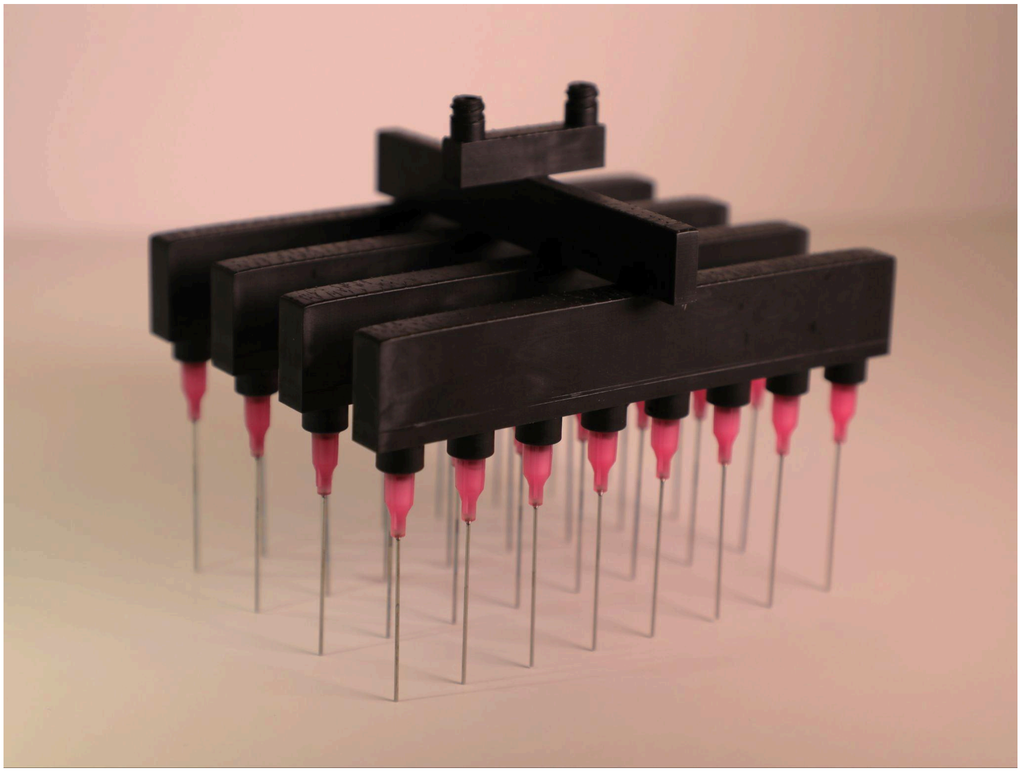

**Figure S6. 32-Multinozzle Printhead.**

| Publication Title | Cost | Scratch or Modification |
|---|---|---|
| Development and characterization of a low-cost 3D bioprinter[1] | **Total $1370** | From scratch |
| Development of a high-performance open-source 3D bioprinter[2] | Custom Parts and FlashForge Finder - **Total $900** | Modification |
| Ultra-Low-Cost 3D Bioprinting: Modification and Application of an Off-the-Shelf Desktop 3D-Printer for Biofabrication[3] | $169 in parts + $200 Anet A8 Desktop 3D Printer Prusa i3 DIY Kit - **Total $369** | Modification |
| A Custom Ultra-Low-Cost 3D Bioprinter Supports Cell Growth and Differentiation[4] | Parts + Anet A8 - **Total $230** | Modification |
| Accessible bioprinting: adaptation of a low-cost 3D-printer for precise cell placement and stem cell differentiation[5] | $200 in parts + $1000 Felix 3.0 - **Total $1200** | Modification |
| Design and implementation of a low cost bio-printer modification, allowing for switching between plastic and gel extrusion[6] | $300 in hardware costs + $900 Prusa I3 MK3 3D - **Total $1200** | Modification |

| | | |
|---|---|---|
| A super low-cost bioprinter based on DVD-drive components and a raspberry pi as controller[7] | **Total $210** | From Scratch |
| Feasibility of Bioprinting with a Modified Desktop 3D Printer[8] | Modification costs (unreported) + MakerBot Replicator 2X Experimental 3D Printer ($2500) - **Total $2500+** | Modification |
| Nydus One Syringe Extruder (NOSE): A Prusa i3 3D printer conversion for bioprinting applications utilizing the FRESH-method[9] | €80 modification + $900 Prusa i3 3D printer - **Total ~$1000** | Modification |
| A Multifunctional 3D Bioprinting System for Construction of Complex Tissue Structure Scaffolds: Design and Application[10] | Not Reported | From Scratch |
| Deployable extrusion bioprinting of compartmental tumoroids with cancer associated fibroblasts for immune cell interactions[11] | **Total £900 (~$1,200)** | From Scratch |
| Bioprinted chitosan-gelatin thermosensitive hydrogels using an inexpensive 3D printer[12] | **Total $1000** | Modification |

**Table S1. Existing low-cost bioprinters in the literature.** Included are their associated costs and whether they are created from scratch or by modifying a commercial off-the-shelf 3D printer.

| Component | Quantity | Cost per unit | Total cost | Source of materials | Product/Model number |
|---|---|---|---|---|---|
| RUMBA+ board | 1 | $23.84 | $23.84 | Aliexpress | 3256804594163368 |
| TMC2208 Stepper Motor Drivers | 6 | $4.4 | $26.38 | Amazon | ASIN: B07RK28VWC |
| 100mm Linear Stepper Motors | 4 | $28.9 | $112 | Alibaba | Model Number: HPB006 -100MM |
| 50mm Linear Stepper Motors | 2 | $17.80 | $35.6 | Alibaba | Model Number: T0601-50MM |
| Power Cord | 1 | $9.99 | $9.99 | Amazon | ASIN: B0D2CBVVLN |
| 12mm M3 Screws | 100 | $0.11 | $11.29 | McMaster | 91290A117 |
| 60mm M3 Screws | 8 | $0.41 | $10.35 | McMaster | 91290A181 |
| M3 Nuts | 50 | $0.07 | $3.75 | Amazon | ASIN: B06XPFLNBS |
| Adhesive Rubber Pads | ¼ of Pack | $1.5 | $1.5 | Amazon | ASIN: B08HN5R9Y3 |
| ¼" 12 by 12 Black Acrylic | 2 | $7.5 | $14.99 | Amazon | ASIN: B07X8F9X4K |
| Syringe dispensing tips | as needed | | | Nordson dispensing tips are used in this paper | |
| Syringes | as needed | | | BD syringes are used in this paper | |

**Table S2: Bill of Materials**. Total printer cost is approximately $250, not including shipping and tax costs. Please note, costs may change and cheaper alternatives may be available.

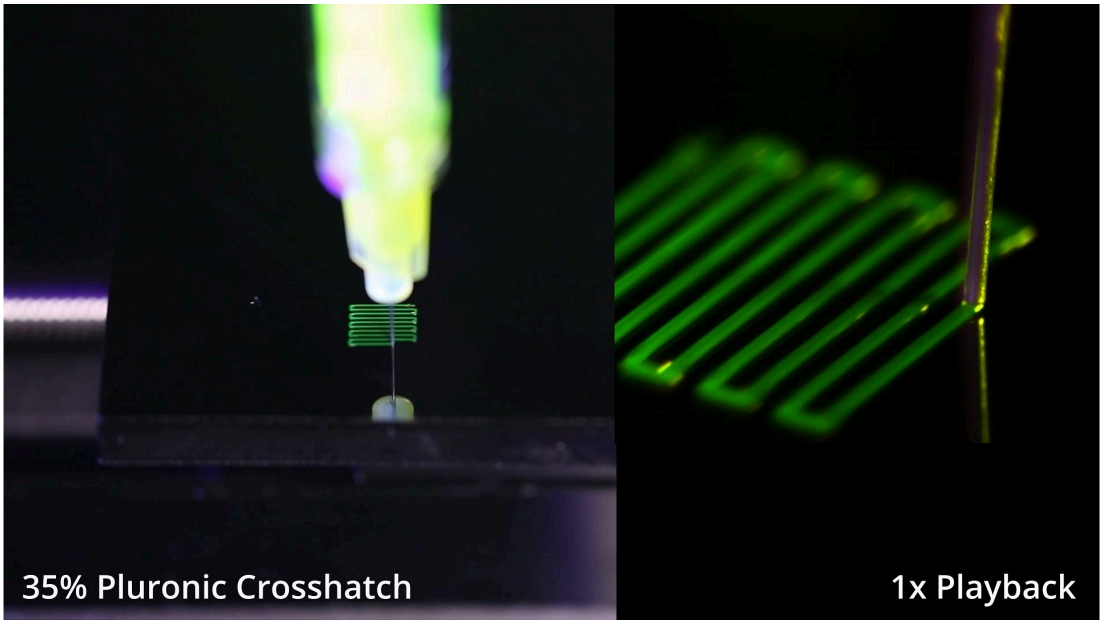

**Supporting Video S1: Multimaterial Crosshatch Print.**

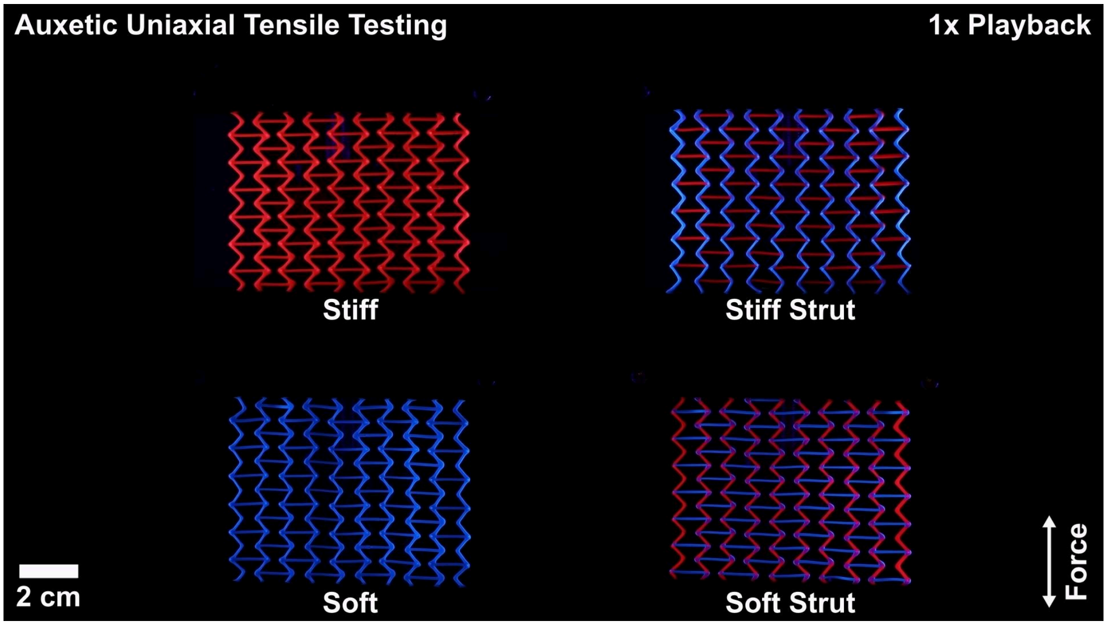

**Supporting Video S2: Auxetic Uniaxial Tensile Testing.**

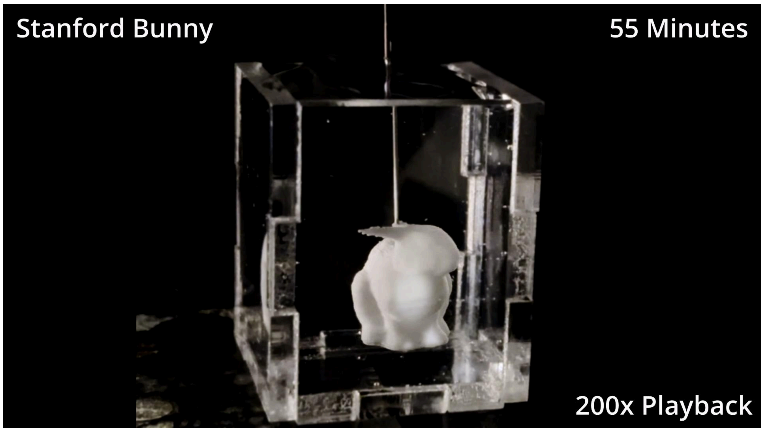

**Supporting Video S3: Embedded Stanford Bunny Print.**

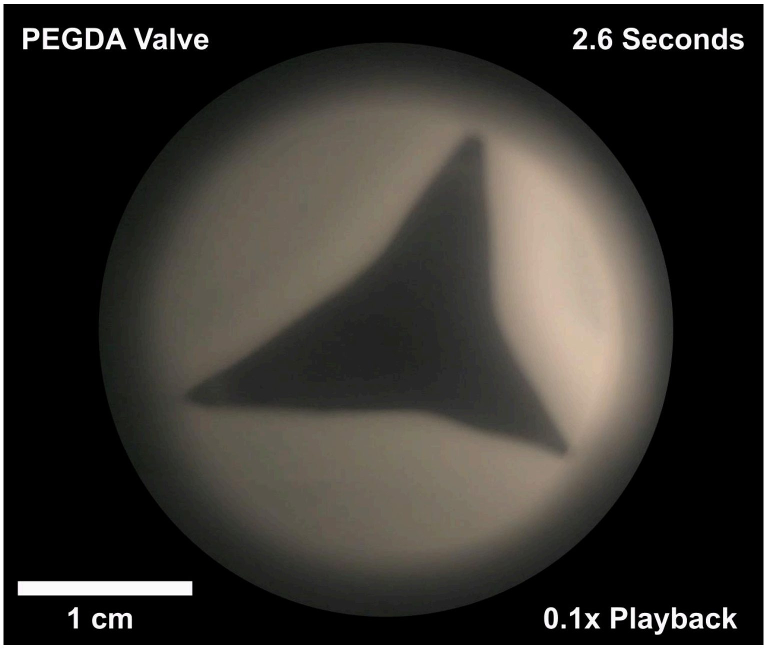

**Supporting Video S4: High-Speed Valve Hemodynamic Testing.** Stroke volume = 60 mL. Heart rate = 60 beats per minute.

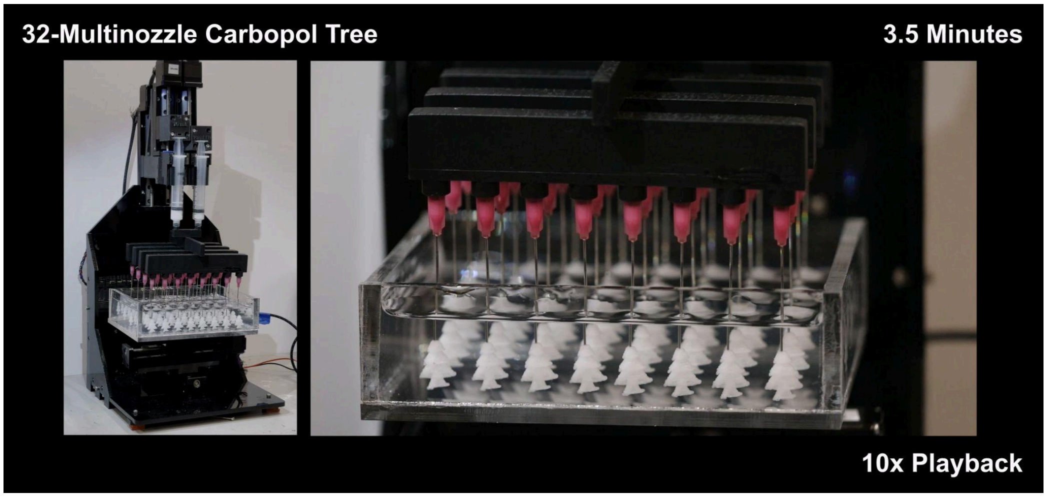

**Supporting Video S5: 32-Multinozzle Tree Print.**

### Supporting Methods: Printess Manual

**Safety:** We remind readers that (1) the Printess is not a toy and contains moving mechanical parts and electronic components, (2) its operation should be closely supervised by someone familiar with standard machine shop precautions, including removing loose jewelry and clothing, and tying hair back, and (3) the motors should be current limited using the driver trim-pots at a level that is just sufficient for printer operation.

To build the Printess, you will first need to 3D print and laser cut all the components and assemble the printer, including wiring the RUMBA control board (See “Assembly Instructions”). You will then need to upload the Marlin firmware to the RUMBA board (See “Install Arduino and Upload Firmware”). Subsequently, you can install the Pronterface GUI (See “Install Pronterface”) and set up the Python algorithm to calculate extrusion distances for your G-code scripts (See “Install Python” and “Using Python Script to Calculate Extruder Distances”). If you are unfamiliar with G-code, the “Crucial G-code Commands and Examples” section may be helpful. We have additionally provided a troubleshooting section. If you wish to build any of the additional components, such as the syringe cooler, tissue culture cooling plate, or tip-tilt platform, you will find sections with those instructions following the troubleshooting section.

#### Assembly Instructions

All CAD files, firmware, and scripts are available open-source on Zenodo at https://doi.org/10.5281/zenodo.13173619. The latest versions of these files and the construction and operation manual can be found on Github at:
https://github.com/weiss-jonathan/Printess-Low-Cost-3D-Printer.

Begin by 3D printing the following CAD files and laser cutting the SVG files for the back and base. The required and optional files are provided in both .stl and .step format within their respective folders. We have had success printing the CAD files on the Mark 2 3D printer (MarkForged, Watertown, MA) using black onyx and carbon fiber materials. Subsequent printer iterations have been successfully printed on a Prusa printer in PLA. For the base and back, ¼” thick 12” by 12” black acrylic sheets were laser cut on a Glowforge machine, but can be cut on any laser cutter accommodating acrylic material.

| **Part name** | **File type** | **Quantity** | **Filename** |
|---|---|---|---|
| Required components | | | **.stl and .step provided** |
| Right support 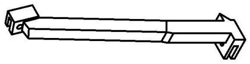 | CAD | 1 | Right Brace Thick.stl<br>Right Brace Thick.step |
| Left support 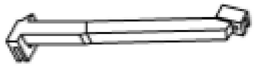 | CAD | 1 | Left Brace Thick.stl<br>Left Brace Thick.step |
| Corner support 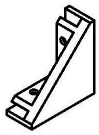 | CAD | 2 | Triangle Brace.stl<br>Triangle Brace.step |
| Base wedge support 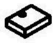 | CAD | 2 | Base wedge.stl<br>Base wedge.step |
| X and Y axis connector 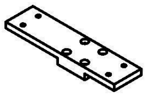 | CAD | 1 | XY Axes Connector.stl<br>XY Axes Connector.step |
| Z axis to extruder connector 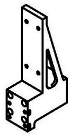 | CAD | 2 | Z-Axes Holder.stl<br>Z-Axes Holder.step |
| *Syringe plunger holder: 1mL 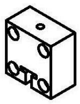 | CAD | 2 | 1mL plunger holder.stl<br>1mL plunger holder.step |

| | | | |
|---|---|---|---|
| *Syringe barrel holder: 1mL 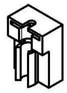 | CAD | 2 | 1mL Barrel Holder.stl<br>1mL Barrel Holder.step |
| Platform attachment: flat 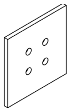 | CAD | 1 | Stage.stl<br>Stage.step |
| Base | PDF | 1 | Acrylic Bottom.PDF |
| Back | PDF | 1 | Acrylic Back.PDF |
| <u>Optional components</u> | | | |
| Syringe plunger holder: 3mL | CAD | optional | 3mL plunger holder.stl<br>3mL plunger holder.step |
| Syringe barrel holder: 3mL | CAD | optional | 3mL Barrel Holder.stl<br>3mL Barrel Holder.step |
| Syringe plunger holder: 5mL | CAD | optional | 5mL plunger holder.stl<br>5mL plunger holder.step |
| Syringe barrel holder: 5mL | CAD | optional | 5mL Barrel HolderL.stl<br>5mL Barrel Holder.step |
| Syringe plunger holder: 10mL | CAD | optional | 10mL Plunger Holder.stl<br>10mL Plunger Holder.step |
| Syringe barrel holder: 10mL | CAD | optional | 10mL Syringe Barrel Holder.stl<br>10mL Syringe Barrel Holder.step |
| Platform attachment: 90mm petri dish | CAD | optional | 90mm Petri Dish Holder.stl<br>90mm Petri Dish Holder.step |
| Platform attachment: 35mm petri dish | CAD | optional | 35mm Petri Dish Holder.stl<br>35mm Petri Dish Holder.step |
| Platform attachment: cube holder | CAD | optional | Cubic container holder.stl<br>Cubic container holder.step |
| Platform attachment: 6 well-plate | CAD | optional | 6 well plate holder.stl<br>6 well plate holder.step |
| Mixing Nozzle | CAD | optional | Mixing Nozzle Body.step<br>Mixing Nozzle Body.stl<br>Impeller.step<br>Impeller.stl<br>Motor Spacer.step<br>Motor Spacer.stl |
| 8-Multinozzle | CAD | optional | 8x1 Bimaterial Multinozzle Printhead.stl<br>8x1 Bimaterial Multinozzle Printhead.step<br>1mL Multinozzle Barrel Holder.stl<br>1mL Multinozzle Barrel Holder.step<br>3mL Multinozzle Barrel Holder.stl<br>3mL Multinozzle Barrel Holder.step |
| 32-Multinozzle | CAD | optional | 8x4 Bimaterial Multinozzle Printhead.stl<br>8x4 Bimaterial Multinozzle Printhead.step |
| Tip tilt platform | CAD | optional | Tip-Tilt Platform.stl<br>Tip-Tilt Platform.step |
| Syringe cooling sleeve | CAD | optional | Cooling Sleeve for 1mL BD syringe.stl<br>Cooling Sleeve - Cooling Sleeve Disc For Vacuum.stl<br>Cooling Plate Vertical - 3D Printed.stl<br>Cooling Sleeve for 1mL BD syringe.step |

| | | | |
|---|---|---|---|
| | | | Cooling Sleeve - Cooling Sleeve Disc For Vacuum.step<br>Cooling Plate Vertical - 3D Printed.step |
| Cooling bed | CAD | optional | Laser Cut from 3mm Acrylic - Cooling Plate Padding.PDF<br>Laser Cut from 6mm Acrylic - Cooling Plate Main Body and Crossbeam.PDF |
| Motor safety stopper | CAD | optional | Motor Safety Stopper.stl<br>Motor Safety Stopper.step |

Description of 3D printed components:

- Supports: The right, left, and corner supports offer stability to the printer and help minimize vibrations during printing. The square base supports are fitted with rubber adhesive pads to lift the printer up slightly from the table surface.
- Axes connectors: These parts allow for the connection between linear stepper motors.
- Platform attachments: A variety of compatible build platforms are available including petri dish, cube holder, and 6 well-plate.
- *Syringe plunger and barrel holders: A variety of syringe mounting designs are available that can accommodate 1mL, 3mL, 5mL Nordson plastic syringes. You can use different sizes on each of the two Z axes, but the plunger and barrel holder on one side must match in size.

To assemble the Printess, follow the guide below using M3 screws. Before beginning, make sure to heat-set square M3 nuts into the cavities in the 3D printed components. Heat-setting requires a soldering iron. Instructions on heat setting can be found here: [https://markforged.com/resources/blog/heat-set-inserts#:~:text=Line%20the%20tip%20of%20the,and%20not%20at%20an%20angle](https://markforged.com/resources/blog/heat-set-inserts#:~:text=Line%20the%20tip%20of%20the,and%20not%20at%20an%20angle).

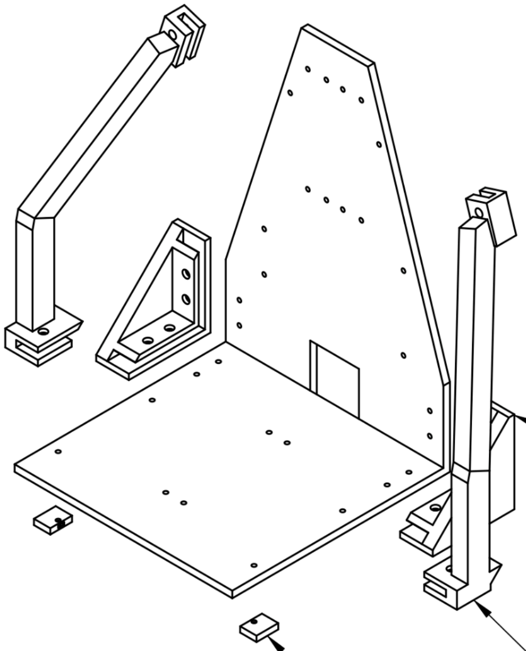

SCALE 1:3

The printer structure should look like this after steps 1 - 3.

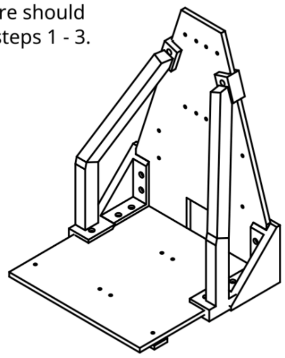

**Step 1)** Attach the left and right holders to the back plate. Repeat for the bottom plate. (M3 x 12 mm screws)

Note: Make sure the back plate is facing the right direction. The side for the Rumba board should be facing the back and should have the label "Rumba Board Here."

**Step 2)** Attach the left and right braces to back and bottom plates.
(M3 x 14 mm **black** screws)

**Step 3)** Attach the base wedges to the bottom board.
(M3 x 12mm screws)

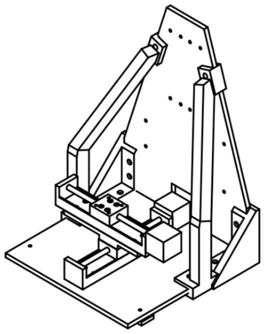

SCALE 1:5

**Step 7)** Guide the wires from both motors through the opening of the back plate.

**Step 6)** Attach another stepper motor with 100mm travel length to the XY connector. Use (M3 x 12mm screws).

Note: There are square nuts embedded into the XY connector, so no need for Hex nuts. This is going to be the X-Axis Stepper Motor

**Step 5)** Attach the XY connector to the stage of the Y-Axis Stepper Motor with the short end facing the right side. (M3 x 12mm screws)

**Step 4)** Attach one of the stepper motors with 100mm travel length to the bottom acrylic plate with the motor side facing towards the back plate. Use (M3 x 12mm and hex nuts with washers) for the attachments.

Note: This is going to be the Y-Axis Stepper Motor.

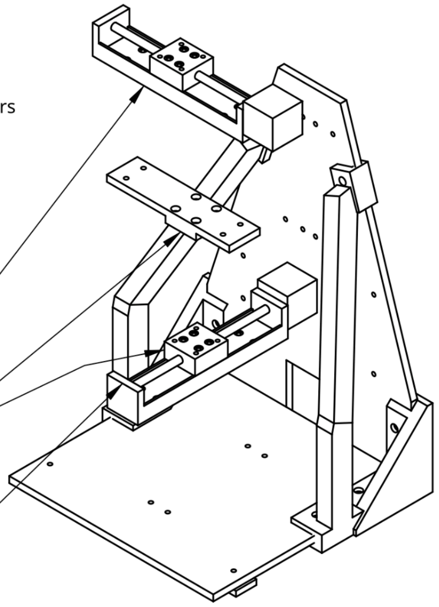

SCALE 1:3

**Step 8)** Attach one 100mm travel length stepper motor to the left side of the back plate. Repeat for the right side.

Use (M3 x 12mm screws and hex nuts with washers) for the attachments

Note: The left stepper motor is the Z-Axis and right motor is the A-axis

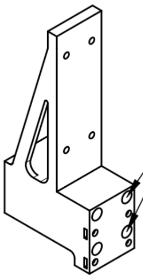

SCALE 1:2

**Step 9)** Attach the Z and A axis holder to the stages of their respective stepper motors.

Use (M3 x 60mm screws) for the attachments. The screw goes through these holes as shown

**Step 10)** Attach one 50mm travel stepper motor to the axis holder. Repeat for the other axis holder.

Use (M3 x 12mm screws)

Note: There should be square nuts in the back of the axis holder.

Of these smaller stepper motors, the left is the B-axis and the right is the C-axis.

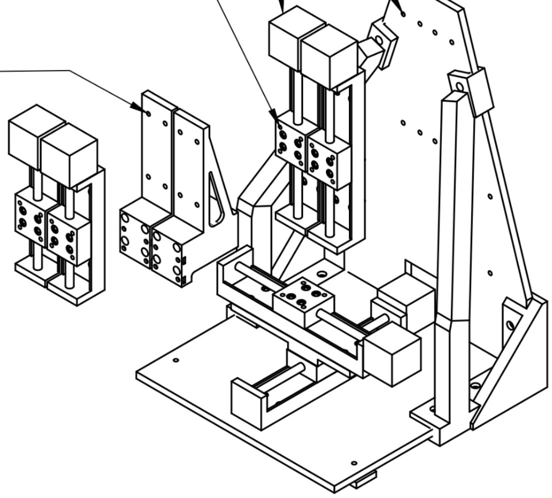

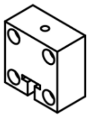

**Step 11)** Attach one of the plunger holders to the stage of the B-axis. Repeat for the C-axis.

Use (M3 x 12mm screws) for the attachments.

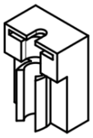

**Step 12)** Attach one barrel holder to the Z-axis connector. Repeat for the A-axis holder.

Use (M3 x 12mm screws) for the attachments.

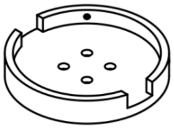

**Step 13)** Attach the print platform to the stage of the X-axis step motor.

Use (M3 x 12mm screws) for the attachments.

Note: The print platform can vary in shape and size.

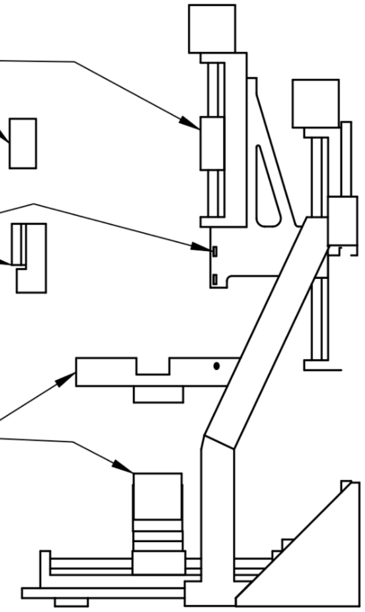

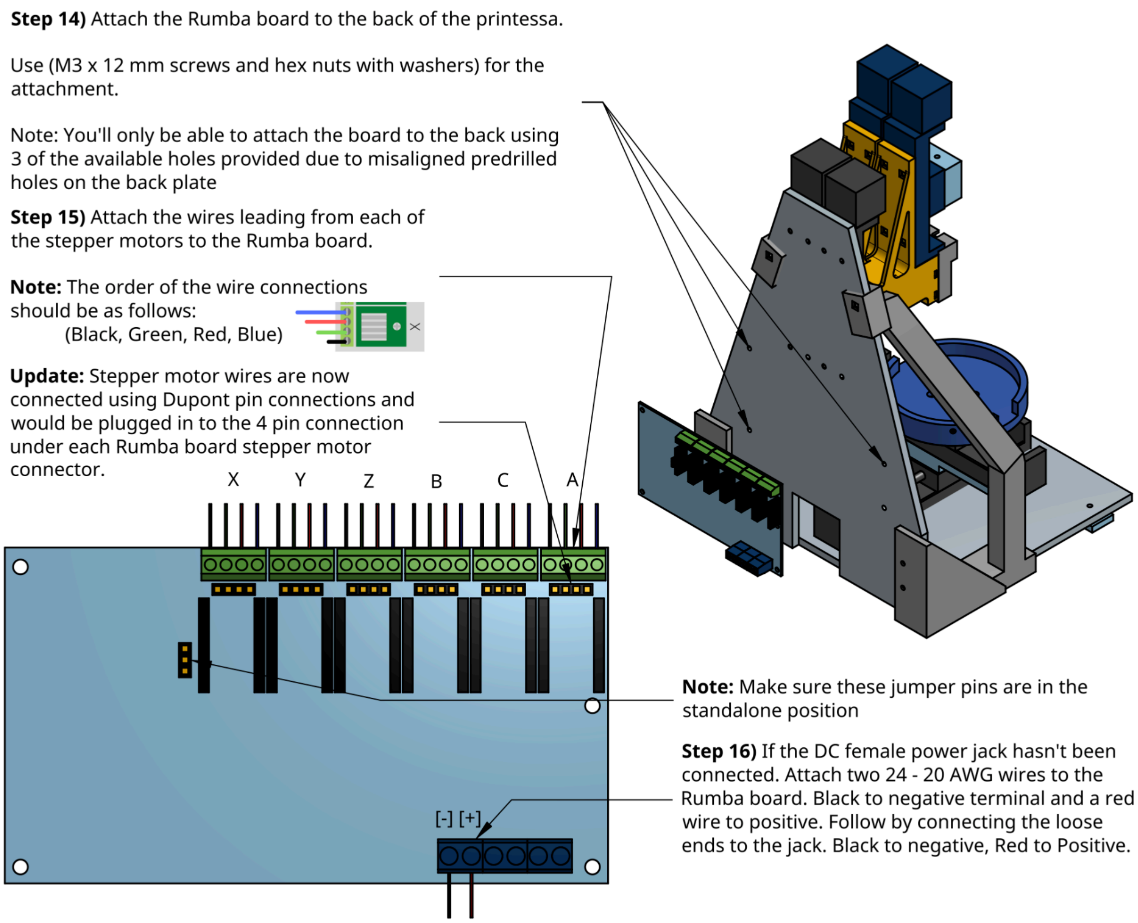

Dupont pin connectors as mentioned above are optional and can make the stepper motor connection more secure and easy to disassemble if necessary. Video on how to use Dupont pin connectors.

| Component | Cost | Source of materials | Product/Model number | Notes |
|---|---|---|---|---|
| 4-Pin Dupont Connector Kit | $8.99 | Amazon | ASIN: B07BDJ63CP | If you don't have a crimping tool, there are other kits available that include one. |

Step 17) Make sure the Power Select Pin is set to Standalone power.

Step 18) Plug in the power adapter and attach the power jack to the RUMBA board and use the multimeter to confirm if the voltage is acceptable by putting the negative common probe to the negative terminal and red probe to the positive terminal of the female jack.

Step 19) To check if the stepper motor driver is correctly tuned. Using the multimeter, have the negative probe touching the negative terminal of the RUMBA board and positive probe touching the potentiometer of each of the stepper motors. The acceptable range is 0.36V to 0.4V. After this, assembly is complete.

After completing building the printer, you will need to connect a computer to the RUMBA board mounted on the back of the printer via the included mini USB cord. You can then proceed to 1) uploading the Marlin firmware to the RUMBA 2) installing the Pronterface GUI and 3) setting up the python script to calculate extrusion distances for your G-code.

**Install VScode/Arduino and Upload Firmware**
[https://www.arduino.cc/en/software](https://www.arduino.cc/en/software)

Arduino is used to upload the firmware to the RUMBA Board. Note: [VSCode](VSCode) has a plugin called Auto Build Marlin that compiles and uploads firmware 10x faster than Arduino does:

Install plugin:

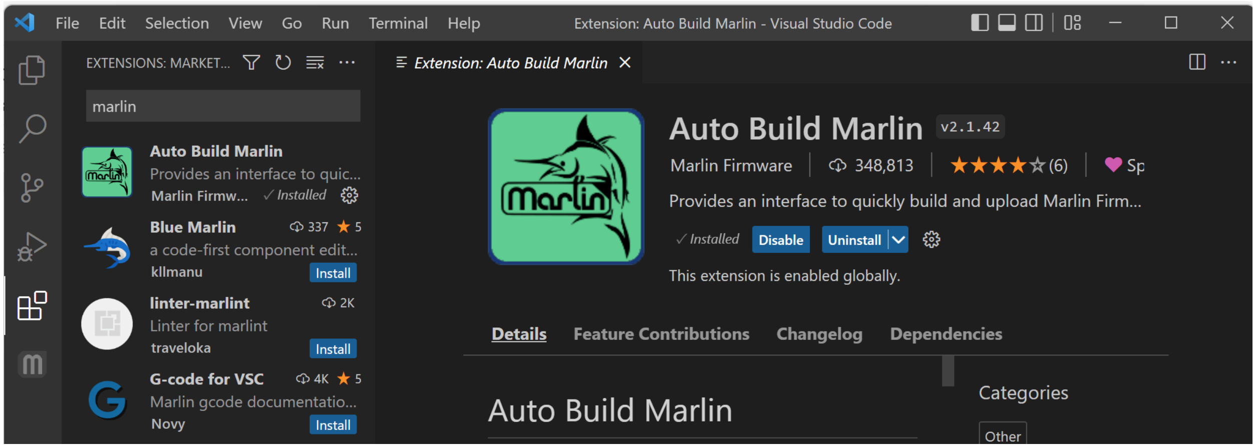

Select Marlin Folder:

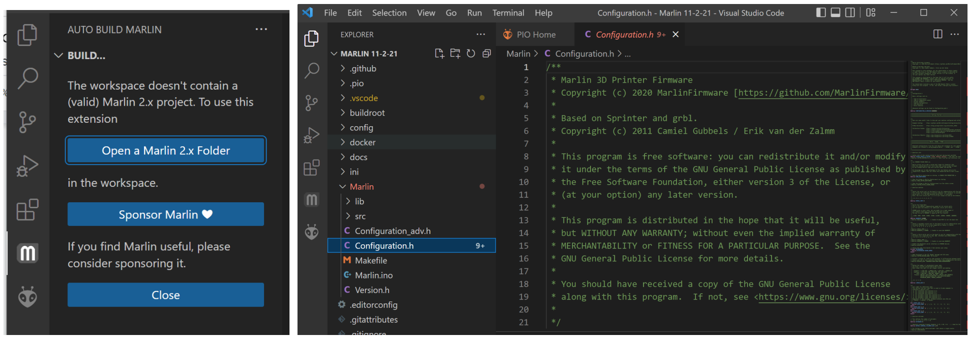

Show ABM Panel:

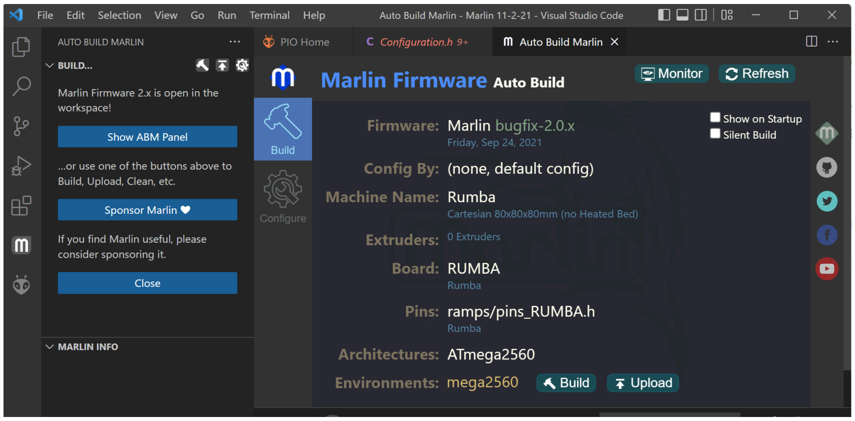

**Do not have the printer connected via Pronterface while uploading firmware.**

If you would rather use Arduino:

1. Download the Marlin Firmware file provided in Zenodo. From within that folder, you can use the version for RUMBA or RUMBA32 depending on which board you have.
2. In the Marlin subfolder, open Marlin.ino in Arduino
3. Ensure that the RUMBA is plugged in and that the lights are on. Connect the board to the computer using a mini USB cable.
4. In Arduino, go to Tools -> Port and select the port that your board is connected to (mine is “usbmodem…”). If you aren’t sure which port it is connected to, disconnect and reconnect, and look at which option disappears and reappears.
5. Tools -> Board and select “Arduino Mega or Mega 2560”
6. Ensure Tools -> Processor is “ATmega2560”

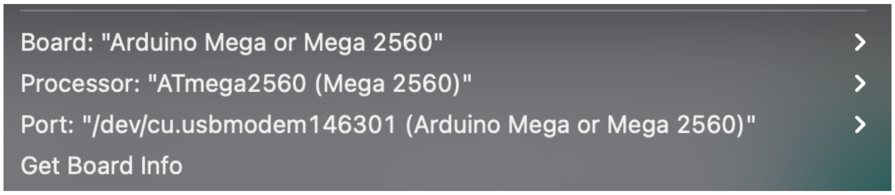

7. Click upload above the Marlin tab (Right arrow). There should be a message saying “Compiling Sketch” below. This will take 10-15 minutes. Ensure your laptop is plugged in as this is an energy intensive process.

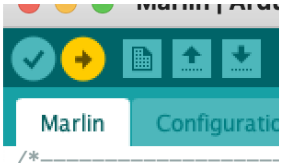

**Install Pronterface**

Precompiled version to download the app:
[https://github.com/kliment/Printrun/releases](https://github.com/kliment/Printrun/releases)

Download the appropriate file for your operating system as shown below:

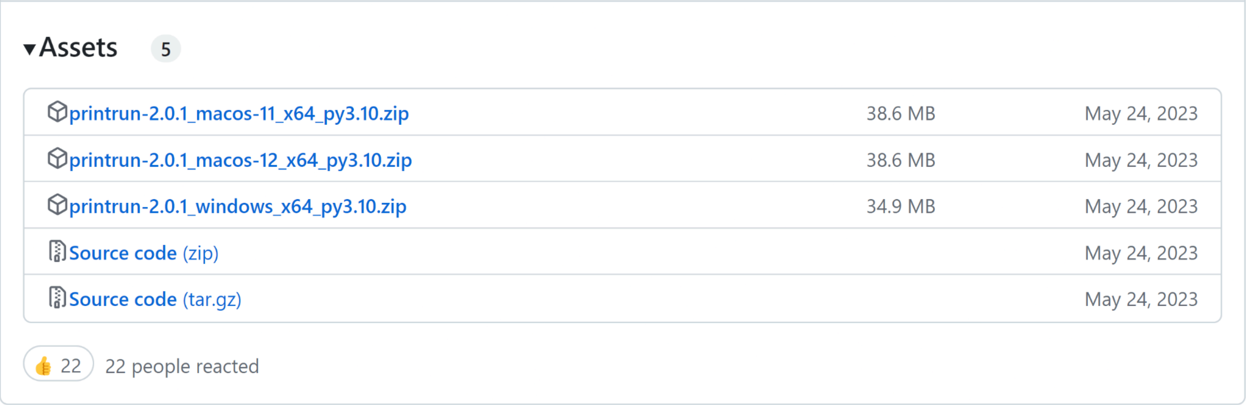

X64 is 64 bit (try this one first)
X86 is 32 bit

**Pronterface Guidelines**

- Connect your printer to your computer, click "Port" to refresh the available inputs, then click the drop down next to it and select the port your printer is connected to.
- Ensure the baudrate is "@115200"
  - If you see errors that include "line number" or "checksum," reduce the baudrate
- Change XY motor rate to 300. It is default to 3000. This is too fast for the motors and they will stall:

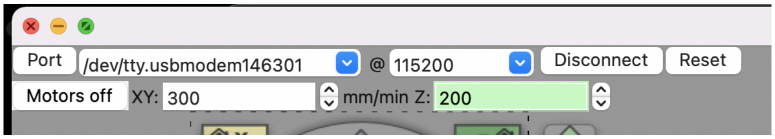

- In general, use the console in the bottom right to send lines of GCode to the printer:

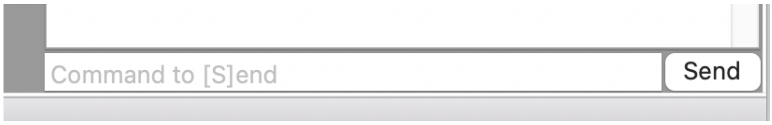

- You may use the GUI controller to manually move the X, Y, and Z axes for quick adjustments, but **BEWARE** using these buttons will automatically reset your positioning system to G90 (see [GCode](#) below) even if you set it to G91 in the console. During experiments, we strongly suggest not using the GUI controller at all. You can move axes by sending commands such as "G1 X10" to the console. If you use the GUI, get into the habit of calling G91 before any motion commands in the console.

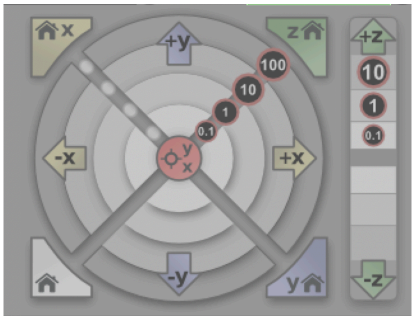

- To upload .txt GCode programs to run, click “Load file” at the top center. “SD, Print, Pause, Off” don’t do anything, so do not click those.
    - When uploading GCode text files to Pronterface, use .txt plain text, NOT rich text format (RTF). If using TextEdit on mac, go to preferences and select “Plain Text,” then make a new document.
- If you need to emergency stop the printer in the middle of a program, you have to either click “Disconnect” FOLLOWED BY “Connect” or unplug the printer from the outlet. Note that every time you reconnect the printer, you need to reset calibrations (See G92 in GCode section below). This is not very convenient, sorry!

Optional:

Making Macros in Pronterface:
[https://github.com/kliment/Printrun/blob/master/README.md#using-macros-and-custom-buttons](https://github.com/kliment/Printrun/blob/master/README.md#using-macros-and-custom-buttons)

**Install Python**

To check current python version in terminal (mac) or command prompt (windows), type the below command:

python --version

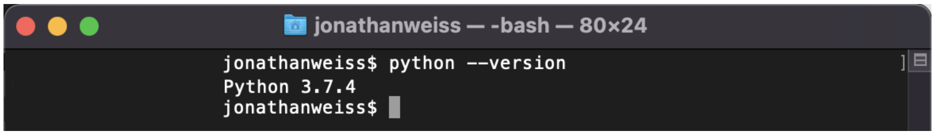

If you do not have Python 3 and need to download python, do so here:
https://www.python.org/downloads/

If you use Windows, make sure python is added to your environmental variables. The complete path of python.exe can be added by:

1. Right-clicking This PC and going to Properties.
2. Clicking on the Advanced system settings in the menu on the left.
3. Clicking on the Environment Variables button on the bottom right.
4. In the System variables section, selecting the Path variable and clicking on Edit. The next screen will show all the directories that are currently a part of the PATH variable.
5. Clicking on New and entering Python's install directory.
    a. C:\Users\...\Python

After downloading python 3, if your python --version still says you have Python 2, you can force python 3 to run by calling "python3" instead of "python". For more permanent solution, check out the forums below:

Windows:
https://medium.com/@ryanmillerc/install-python-3-in-locally-in-appdata-alongside-python-2-in-windows-10-fe4287708429

Mac:
https://stackoverflow.com/questions/43354382/how-to-switch-python-versions-in-terminal

It is easy to run python scripts using your terminal. We will need to run a python script to calculate extrusion distances. More on this later.

**Crucial GCode Commands and Examples**

Familiarize yourself with all of the axes first. Stage Axes: X, Y; Vertical Axes: Z, A; Extrusion Axes: B, C.  Practice moving each axis to learn which axes correspond to which motors and their polarity: for the vertical axes, positive motion is upwards, while for the extrusion axes, positive motion is downward.

Refer to https://reprap.org/wiki/G-code for more information, commands, examples, etc.

<table>
<tr><th>Command</th><th>Description</th></tr>
<tr><td>G90</td><td>Set system to Absolute Positioning. When using absolute positioning, all coordinates you provide will be read in the coordinate system you define. Thus, you need to first define your coordinate system, usually by designating an origin (0, 0, 0). See G92 below. Run on its own line.</td></tr>
<tr><td>G91</td><td>Set system to Incremental Positioning. All distances will be relative to the current location. Run on its own line.</td></tr>
<tr><td>G92</td><td>Recalibrate current position. Use mainly to set current position to origin.<br><br>G90, G91, G92 Example (refer to diagram below). Our goal is to move our location from the origin in the bottom left to points 1, 2, and then 3:<br><br>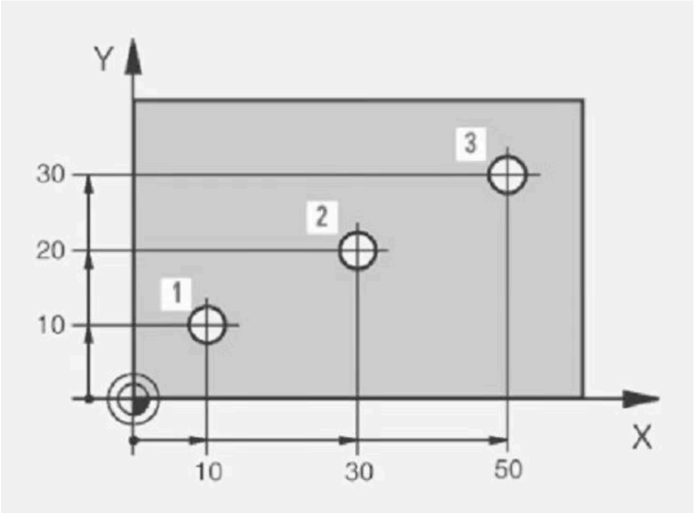
<br><br>G90; Set system to absolute positioning<br>G92 X0 Y0; Set current positions in (X, Y) to (0, 0)<br><br>G1 X10 Y10 F200; Move to the coordinates (10, 10) in (X, Y). This is point 1.<br>G1 X30 Y20 F200; Move to the coordinates (30, 20) in (X, Y). This is point 2.<br><br>G91; Set system to incremental positioning</td></tr>
</table>

<table>
<tr><td></td><td>G1 X20 Y10 F200; Move 20 in X and 10 in Y<br><br>With respect to our defined coordinate system above, we are now at (50,30) in (X, Y). This is point 3. Switching to relative to arrive at point 3 was for the sake of example and unnecessary. It is up to you which coordinate system you want to use. You may find one more logical than the other depending on your application.<br><br>Note: You need to recalibrate your position (G92) after reconnecting to the printer every time. In between running scripts, the printer will remember it's absolute location so long as you do not disconnect.<br><br>Another Example GCode to make a 5x5mm square:<br><br>G91; Relative<br>G1 X5 Y0 F300<br>G1 X0 Y5 F300<br>G1 X-5 Y0 F300<br>G1 X0 Y-5 F300<br><br>G90; Absolute<br>G92 X0 Y0; Define current location as (0,0)<br>G1 X5 Y0 F300<br>G1 X5 Y5 F300<br>G1 X0 Y5 F300<br>G1 X0 Y0 F300</td></tr>
<tr><td>G1</td><td>Move in a straight line.<br><br>Example (refer to diagram below):<br>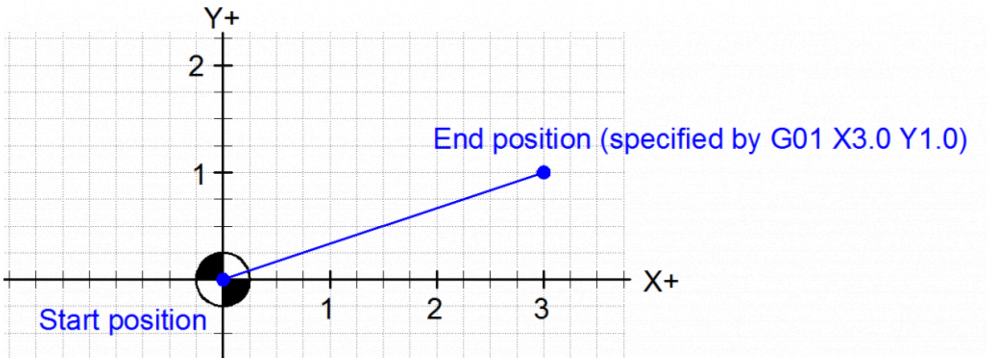<br><br>G90; Absolute Positioning<br>G92 X0 Y0<br><br>G1 X3 Y1 F300<br><br>Note: G1 and G01 are the same. Use G1.</td></tr>
<tr><td>G2, G3</td><td>Clockwise (G2) and counterclockwise (G3) arcs / circles.</td></tr>
</table>

| | |
|---|---|
| | **Example** (refer to diagram below):<br>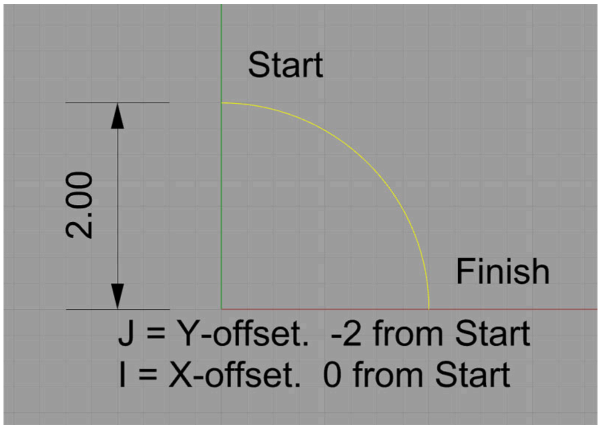<br><br>G91; Incremental Positioning<br>G2 X2 Y-2 I0 J-2<br><br>X and Y indicate the point where circle should end (omitting will default to full circle)<br>I and J indicate the midpoint *relative* to the starting point. (I and J are always read in relative even when you are in absolute positioning).<br><br>Note: If you specify a point (X, Y) that is not on the circle, then the nearest point on the circle will be selected. Please use ncviewer for practicing these commands to understand how they work. |
| F | Define speed of motor for X, Y, Z, and A movements (units in mm/min).<br><br>Use F300 as starting speed. Higher speeds may stall the motors.<br><br>Include an F command on the first motion line you give to the printer after connecting to Pronterface. The printer will remember this speed until you explicitly change it or disconnect, but it does not hurt to include it on every relevant line as we have done in the examples thus far.<br><br>Note: This does NOT influence the speed of the extruders. Pronterface will automatically adjust the speed of the B and C axes to move the distance you specify by the time the X, Y, Z, A translations finish. |
| ; | Anything that comes inline after “;” is a comment and ignored by the computer. |

**Optional Commands**

| Command | Description |
|---|---|

| | |
|---|---|
| G0 | Jog (movement with no extrusion). You do not need this command for our printer as extrusion is controlled explicitly by the extrusion axes. Just **omit B or C** if you want to move with no extrusion. |
| G4 | Dwell. Wait a certain amount of time before running the next line.<br><br>**Example**:<br>G1 X5<br>G4 P2000<br>G1 X5<br><br>Moves 5mm in X, waits 2 seconds (units after "P" are in milliseconds; Can also use "S" for seconds), then moves 5mm again in X. |
| G60 | Save current Position |
| G61 | Return to saved position<br><br>**Example**:<br><br>G60<br>G61 XY F300<br><br>G60 will save all coordinates of the current position, and this code just moves us to the saved X and Y coordinates:<br><br>https://marlinfw.org/docs/gcode/G061.html |
| M114 | Get current position in absolute coordinates. |
| M2 | End code. In general, include this at the end of every Gcode script, but it happens to be unnecessary for Pronterface. |

We have provided an example G-code file in the repository in the "Python Extrusion Distance Script" folder under the filename "Demo_G-code.txt" that you may use to gain familiarity with the printer.

**GCode Visualizer**

- https://ncviewer.com/
- Copy your Gcode into this program to visualize the paths it will make.
- May not recognize axes names beyond X, Y, and Z, so you might have to edit your code if using other axes that your printer has.

**Useful Terminal Commands for Unix Shell (Mac)**

- pwd: print working directory
    - This lets you see what folder you are in.
- ls: list all the contents in your current directory

    - On windows, use dir instead of ls
- cd: change directory (to select a file or enter a new folder within your current folder)
    - i.e. "cd Desktop" to go to Desktop if you are currently in your user folder.
- Type "python pythonScript.py" without quotations to run a python script. Replace pythonScript.py with the name of your python file.
    - Make sure your python script is in the same directory as any files that it needs.
- **Tip:** If you hit tab after typing a few characters, the full item name will automatically populate.

##### Using Python Script to Calculate Extruder Distances

###### Extrusion Distance Calculations

We developed a python script (most up-to-date version [here]()) to calculate the necessary extrusion distance for each line of Gcode. This program is available for use and modification on Github ([https://github.com/weiss-jonathan/Printess-Low-Cost-3D-Printer/blob/main/Python%20Extrusion%20Distance%20Script/gcode_translator_08_10_23.py](https://github.com/weiss-jonathan/Printess-Low-Cost-3D-Printer/blob/main/Python%20Extrusion%20Distance%20Script/gcode_translator_08_10_23.py)) and in the Zotero repository. The program inputs a .txt file of G-code and outputs the file with calculated extrusion distances. It can be run using the terminal or an IDE such as Pycharm. It may be necessary to include a small initial extrusion move to pressurize and syringe prior to printing and a final small reverse extrusion to depressurize after printing. The extrusion calculations rely on the fact that the cylindrical geometric volume that is pushed down in the syringe barrel is equal to that extruded from the nozzle, allowing us to calculate the height, which is the extrusion distance. Extrusion distance can be defined as the distance in millimeters that the stepper motor connected to the syringe plunger moves. Extrusion distance is calculated as follows:

| | |
|---|---|
| **E** = extrusion distance<br><br>**k** = extrusion coefficient, a user determined scaling factor<br><br>$\mathbf{b_d}$ = syringe barrel diameter<br><br>$\mathbf{n_d}$ = nozzle diameter<br><br>**l** = length of path (i.e., G1 X5, l =5) | $Volume\ pushed\ =\ volume\ extruded$<br><br>$E * \pi * (\frac{b_d}{2})^2 = k * l * \pi * (\frac{n_d}{2})^2$<br><br>$E = \frac{k*l*(\frac{n_d}{2})^2}{(\frac{b_d}{2})^2}$<br><br>$E = \frac{k*l*n_d^2}{b_d^2}$ |

The extrusion coefficient value, "k," may be changed throughout the code by writing an overriding line (K = new coefficient #, k = #, KK = # or kk = #).

In a simpler explanation, the purpose of this program is that we need to calculate the distance that the extruder axis has to push the plunger in order to extrude a filament of desired volume and diameter. Imagine moving the plunger a certain distance downward while X moves by 5mm, and imagine moving the plunger the same distance while X moves by 10mm. In the latter case, the same amount of material is extruded across a farther distance, so by conservation of mass, the filament will be thinner. We have provided a Python script that will calculate what the extruder coordinate (B and C axes) has to be given the X, Y, Z, and/or A coordinates to extrude a filament of desired thickness.

1. Begin by creating a .txt file and name it “gcode.txt”. This name needs to be exact. On a Mac, you can use the application TextEdit. Paste this code at the top and alter the positioning system and syringe parameters based on your subsequent GCode:

   G91
   Z_syringe_diameter = 4.6
   A_syringe_diameter = 4.6
   Z_nozzle_diameter = 0.2
   A_nozzle_diameter = 0.2
   extrusion_coefficient = 1
   [Add your GCode here]

   Example:

```
G90
Z_syringe_diameter = 4.6
A_syringe_diameter = 4.6
Z_nozzle_diameter = 0.2
A_nozzle_diameter = 0.2
extrusion_coefficient = 3

G1 X24.0 Y0.0 F300.0
G1 Z2 ; NO E
G1 X0 Y-4 F300 ; NO E
G1 Z0
```

   Important Notes:

   - Make sure there is no empty line at the top of the .txt file. The first line must be G90 or G91, depending on whether you are using an absolute or relative coordinate system.
   - If you have inline comments, make sure there is a space between before the “;”.
     - ie G1 X5 Y5 ;comment
   - If you are troubleshooting and running this script several times, make sure you close the gcode_modified.txt file in between runs to avoid confusion between versions.
   - When the **extrusion coefficient** is set to 1, the extruded filament will have the same diameter as your nozzle. Setting it 2 will double the volume. 3 will triple, etc. It is a simple scaling factor for the extrusion axes.

| BD Syringe Size | Inner Diameter (mm) |
|---|---|
| 1 mL | 4.9 |
| 3 mL | 8.6 |

| | |
|---|---|
| 5 mL | 12.0 |
| 10 mL | 14.4 |

2. Run gcode_translator using your terminal (see above section for useful commands) or an IDE such as Pycharm
   a. If your .txt file is named something other than “gcode.txt” (it is easier to just rename your .txt file), you can go into the python script and alter line 5 with the file name:

      file = open(“gcode.txt”, “r”)

   b. Run the code, ensuring that your .txt file and the python script are in the same directory.
   c. The translator will output the file with the extrusion distances calculated as a new file called “gcode_modified.txt”
   d. Double check that your code looks good, and now upload this modified file to Pronterface
      i. We recommend an initial extrusion to pressurize the syringe prior to printing and a final “unextrusion” to depressurize after printing.

         For example, right before the coded path in your gcode, include the line:

         G1 B0.2 F400

         and right after finishing, include the line:

         G1 B-0.2 F400

         Alter the distance as needed.

**Using CURA to Slice 3D Models to GCode**

Go to the Ultimaker Cura Slicer download page.

Download and install the software version corresponding to the user’s operating system. Note: For Mac users, there are two versions available: x64 and ARM64. “x64” is meant for Intel based chips and “ARM64” is for newer macs using the apple silicon chips. The .dmg file is a disk image and the .pkg is an installation package. Choose either one.

Setting Up Printer Configuration Manually:

1. If the program is a new installation, the application may prompt the user to set up a new printer. But if it does not or gets closed by accident, the user can go to Preferences → Printers (left hand menu) → Add New (button top right).
2. Next select “Non Ultimaker printer.”

3. After that, open the drop-down menu “Add a non-networked printer,” scroll down to the “Custom” category, and select the “Custom FFF printer”. The user can change the name of the machine here or do it later.
4. The next screen should be the machine setting page. On this page, the user can enter the size of the build platform as a rectangular 100 mm x 100 mm x 100 mm.
    a. The origin at center checkbox should be checked and all heated options unchecked since the Printess does not have any heating accessories.
    b. Under G-code flavor select “Marlin”
    c. The current setting of the print head is min X: -100, max X: 100, min Y: -100, max Y:100, and the number of extruders is 1 or 2.
    d. For now, the “Apply Extruder offsets…” should be unchecked.
5. Delete everything in the Start G-code and End G-Code section. These sections are where the users would write their own start and end g-code to be applied to Cura’s g-code output.
6. Under each extruder tab
    a. Enter the desired nozzle diameter into the “Nozzle size” field.
    b. Enter the inner diameter of the syringe into the “Compatible material diameter”
    c. Apply any offset in the “Nozzle offset (X/Y)” fields. Extruder 1 has a 0 offset. If using a separate syringe for secondary material, Extruder 2’s offset from Extruder 1 needs to be manually measured and entered.
    d. Enter 0 for the remaining settings

Exporting Printer Configurations:

1. Go to Help → Show Configuration Folder
2. Save the entire folder or folder contents into a separate folder

Importing Printer Configurations:

1. Go to Help → Show Configuration Folder
2. Copy the entire folder into the existing Cura configuration folder.

Adjusting the Print Settings:

1. There are three tabs in the middle: PREPARE, PREVIEW, MONITOR. The user should only be concerned with the PREPARE and PREVIEW.
2. Under PREPARE, there is a button that prompts the user to open a file and 3 drop down menus. The leftmost drop down menu selects the desired Printer configuration. The middle menu is for activating the desired extruder and material. The last menu is the Print Settings.
3. Under Print Settings, select the “show custom” setting button. This gives the user a plethora of customizations for changing the behavior of the g-code output.
4. When hovering over any of the categories, a slider icon appears on the right, which opens the “setting visibility” menu. Here, the user can access options like line width, extruder retraction distance, and speed. Go through all the options and select the ones you would like to change. It is good to play around with these settings to see what they do to optimize your print. Some useful ones include Wall, Top/Bottom, Infill, and every Print Speed option.

5. In the Print Settings drop down menu, go to the speed category and change all the speeds to 5 mm/sec. Under the Travel category, change the retraction speed to 2 mm/sec. This limits the g-code output to a feed rate of F300. Under Cooling, uncheck the Enable Print Cooling box.
6. When finished, click the blue floppy disk save icon on top. This prompts the user of the changes made and that the settings to be saved under a new name or to overwrite an existing customized setting.
7. To export and import custom Printer Setting profiles, go to Preferences → Configure Cura and in the left-hand side menu select Profiles. Here the user can click the Import button on the top right to import a profile. To export a profile, select the custom profile and under the hamburger button ≡, and select export.

**Troubleshooting**

**I get errors when trying to run the python script**

- Try to get rid of all the inline comments from your file before running it (lines that start with a ; should be ok).
- Make sure you have properly named your .txt file.
- If you have G2 or G3 motion, make sure this is in terms of I and J and not R.
- If you suspect there is a Python version issue, try reinstalling python 3. You can also run python scripts by calling "python3" instead of "python".
    - Example: `% python3 gcode_translator_1_27_21.py`

**I get errors when uploading .txt files to Pronterface, or the program is not running once uploaded.**

- Disconnect and reconnect
- Make sure your file is either a .txt or .gcode. (.rtf files DO NOT work.)

**The material coming out of my syringe does not stick to the substrate (glass slide, petri dish, etc.)**

- Most likely your syringe is too high off the substrate. Before starting your program, make sure the syringe is ~0.2mm above the surface. You can manually move the syringe down so that it just barely touches the surface, and then move up by ~0.2mm. You may need to adjust this value depending on how much material you are extruding (i.e. higher height for higher extrusion coefficients).

**When printing materials on agar gel, the material seems to melt.**

- Pluronic will quickly absorb water from the gel and pool, so it will be difficult to make solid structures. Pluronic can be more useful as a medium through which to deliver other molecules of interest to your bacteria. Avoid embedding bacteria in pluronic as it is a detergent and will harm bacteria with prolonged exposure (edge contact is OK). If you must use pluronic and need it to maintain its structure, try using a very high percentage (45%+). This may slow the water absorption process.
- Alginate exhibits a similar behavior but can be avoided by crosslinking with higher concentrations of calcium (try this first) and/or alginate. It is better to embed bacteria in alginate as it doesn't actively harm bacteria.

**My extrusion doesn't start at the very beginning of the print, and it leaks after the print is finished or during segments where I don't want extrusion.**

- You need to include a pressurization command (i.e. G1 B0.2) whenever starting an extrusion to start the flow of the material out the syringe. You also need to include a depressurization command (i.e. G1 B-0.2) after you are done extruding to stop leaking from happening. You will need to adjust how much to pressurize or depressurize via trial-and-error. B+/-0.2 is a good place to begin.

**My syringe runs into the ground or my axes are generally not moving how I want them to when I start a program.**

- Double check that your GCode is correct.
- 99% of the time this happens, you are either in absolute positioning (G90) and forgot to zero using G92 before starting or **you used the GUI to manually position the axes, which resets the positioning system to G90,** and forgot to switch it back. 1% of the time this happens, it is because your motor wiring came loose.

**When I try to move a motor, it makes a sound but doesn't actually move.**

- You are probably trying to run the motor too quickly. Make sure you include an F command at the end of your motion line. We recommend not moving faster than F300.
    - i.e. G1 X10 Y10 F300
- If the bearing is not lubricated well, the table can get stuck. Try running the motor while pushing with your hand to get the table unstuck. If this works, go back to the original spot to see whether the motor gets stuck again. If it appears that that one spot needs lubrication, then add a little of lubricant (SHC-100) to the spot on the bearing where the table was stuck. Use gloves and rub the lubricant around the affected spot with your hand, then discard glove in trash. Run motor back and forth over the spot to further distribute the lubricant.
- If the issue persists, the driver board may not be supplying the motor with enough current to drive the stepper motor and you may need to adjust the potentiometer on the Marlin board connected to that axes.

**My board keeps crashing with the rapid yellow flashing light.**

- Unplug and replug in the printer. If this keeps happening, check the power supply. If it is rated 36V, that's probably the issue since the RUMBA board is technically only rated up to 35V. Swap with a 12 or 24V 4A power supply.

**Optional Components Assembly**

**Mixing Nozzle Assembly**

Additional required components for Mixing Nozzle

| Component | Quantity | Cost per unit | Total cost | Source of materials | Product/ Model number | Notes |
|---|---|---|---|---|---|---|
| 127 RPM Mini Econ Gear Motor | 1 | $12.99 | $12.99 | ServoCity | 638394 | Can substitute for other motors if desired. |
| Push-to-connect tube fitting | 1 | $3.32 | $3.32 | McMaster | 51115K546 | |
| 20 mm long M3x0.5 mm screw | 2 | $0.0818 | $0.1636 | McMaster | 91292A123 | Longer screws are also okay. |
| 8 mm long M3x0.5 mm screw | 2 | $0.0545 | $0.109 | McMaster | 91292A112 | |
| M3x0.5 square nuts | 2 | $0.1428 | $0.2856 | McMaster | 97258A101 | |
| Epoxy | 1 | $32.04 | $32.04 | McMaster | 7370A38 | Can use alternative adhesive if desired. |

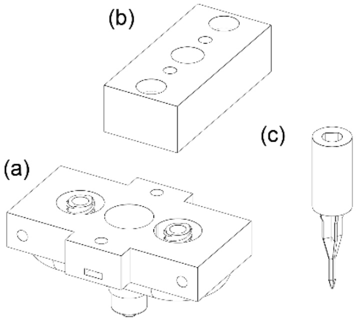

**Step 1)** 3D print the main printhead (a), impeller (b), and motor spacer (c) on a Formlabs 3B+ printer using the BioMed Black resin

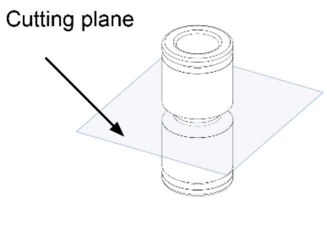

**Step 2)** Cut the push-to-connect tube fitting (McMaster part # 51115K546) at the middle (see cutting plane) and sand the connector until flat

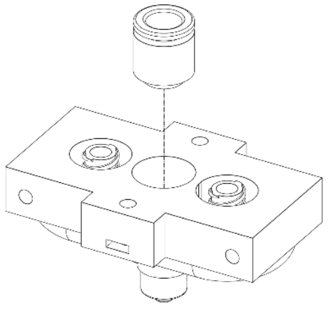

**Step 3)** Adhere the cut push-to-connect fitting into the center slot of the 3D printed main printhead using epoxy. Allow it to cure following the epoxy manual

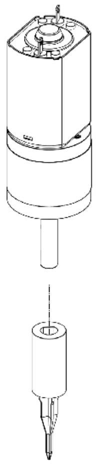

**Step 4)** Press fit the printed impeller onto the D bore shaft of the motor (ServoCity SKU # 638394)

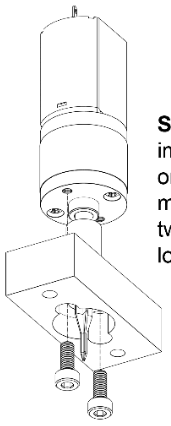

**Step 5)** Screw motor/ impeller assembly onto the 3D printed motor spacer using two M3x0.5mm 8mm long screws

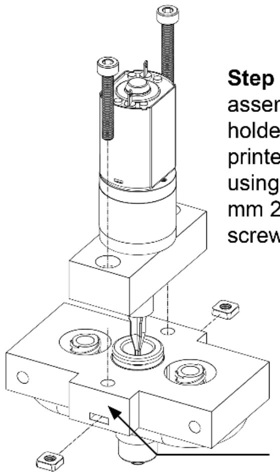

**Step 7)** Screw the assembled motor holder onto the 3D printed extruder using two M3x0.5 mm 20 mm long screws

**Step 6)** On both sides, slide square nuts into the 3D printed extruder

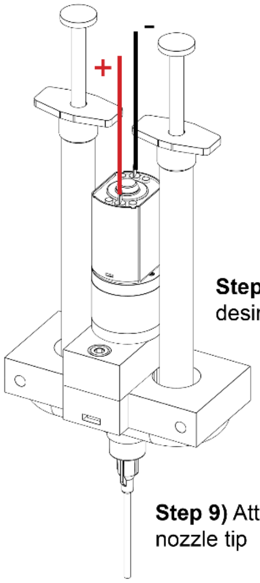

**Step 8)** Attach desired syringes

**Step 9)** Attach desired nozzle tip

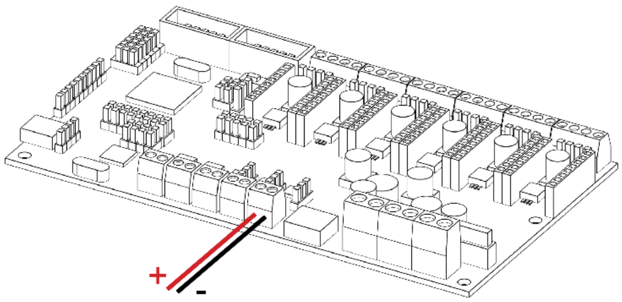

**Step 10)** Connect the DC motor to the fan 0 pin on the Rumba board. Speed can be modulated using the Marlin fan command M106 P0 S[speed], where speed is the duty cycle, which is out of 256 (i.e. S256 would be 100%), and M107 [off].

#### Syringe Cooler Assembly

### 1ml BD Syringe Cooler

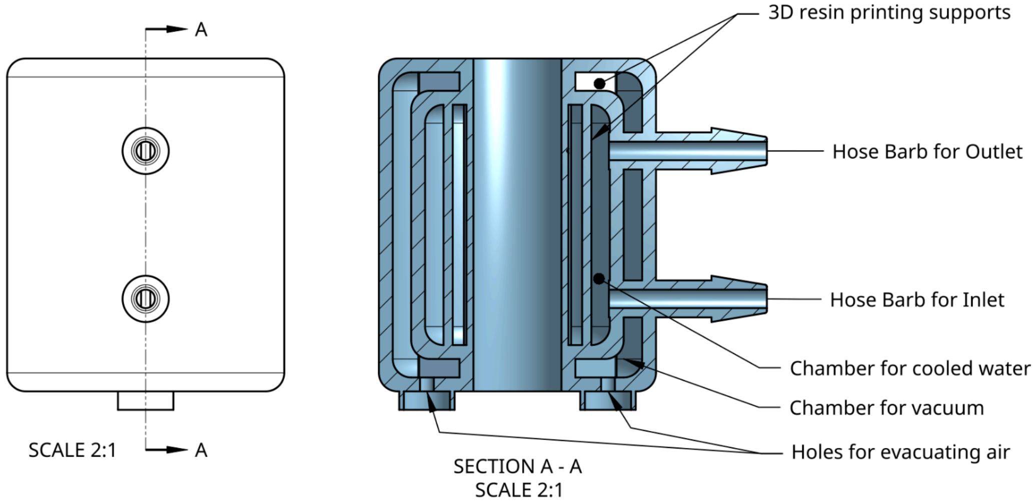

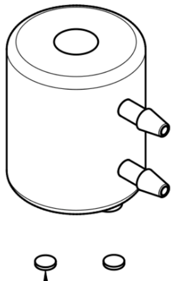

- Syringe cooler was printed using Formlabs Rigid 10k Resin.
- There are two chambers: one for the chilled water and one for the insulating vacuum.
- The vacuum was generated by placing the cured cooler in a vacuum chamber with the holes for evacuating air facing upwards.
- Two small discs were placed on top of the holes with a dab of resin covering them.
- With the vacuum on, the existing air in the chamber forced out and as the air was reintroduced, the external pressue forced the disc shut.
- The dab of resin was then UV cured to seal in the vacuum.
- This design is made for the 1mL BD syringe and secured with a O-ring or hose clamp.

#### Tissue Culture Cooling Plate Assembly

Additional required components for cooling plate

| Component | Quantity | Cost per unit | Total cost | Source of materials | Product/Model number |
|---|---|---|---|---|---|
| Aluminum Water Cooling Block | 1 | $9.5 | $9.5 | Amazon | 757233296242 |

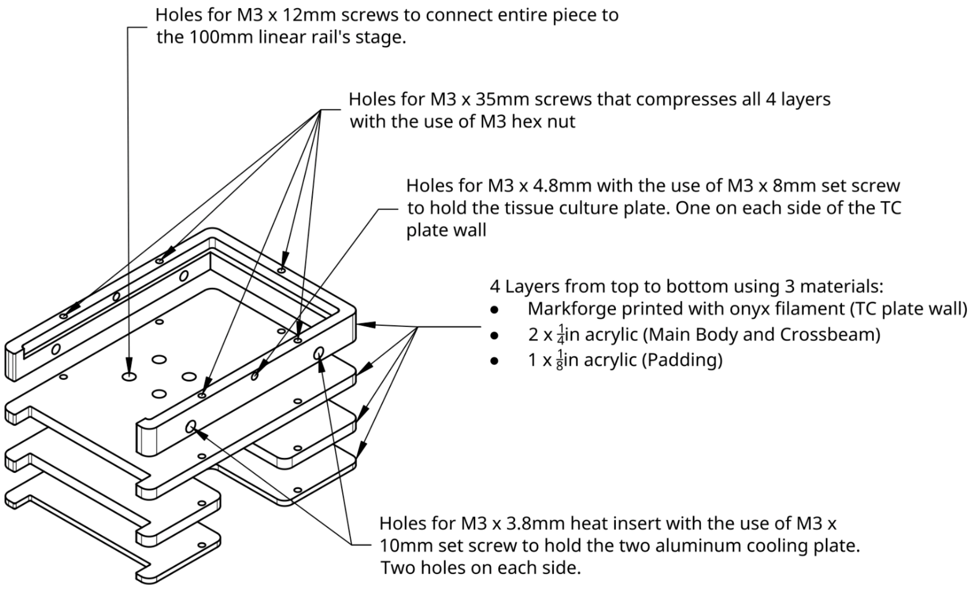

This cooling apparatus pumps chilled DI water from a plastic or glass jar that has two 1/8" inch trough bulkhead barbs attached and the pump is any perisltaic pump that can handle 4mm tubing with 3mm ID. The chilled water first goes through the syringe cooler, then to the aluminum blocks of the plate cooler and returns the DI water to the reservior.

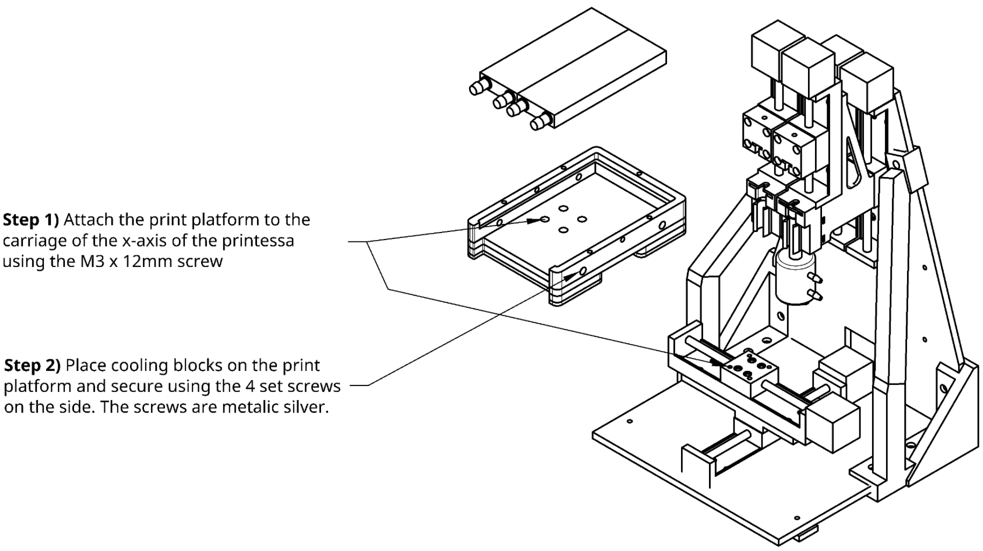

### Cooling Setup

**Step 3)** Slip the BD 1mL syringe into cooling sleeve and place an 8mm O-ring or 8mm hose clamp on the leur lock side of syringe to hold up the cooling sleeve.

**Optional:** Add a "tiny" bit of thermal grease on the syringe to improve contact to the cooling sleeve

**Step 4)** Insert the syringe into the barrel holder and plunger the plunger holder.

**Step 5)** Connect the outer barb of one cooling block to the outer barb of the second block using one of the 7mm ID x 145mm tubes.

**Step 6)** Attach one 7mm ID x 145mm tube to one of the inner barbs. Attach the last 7mm ID x 145mm tube to the last remaining barb. Attach 8mm to 4mm reducer to the open ends of both tubes. One tube is going to going to be the inlet and the other is outlet for this set of cooling blocks.

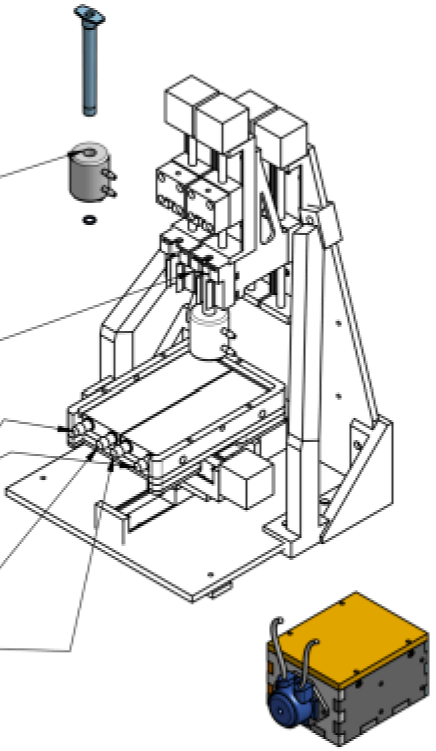

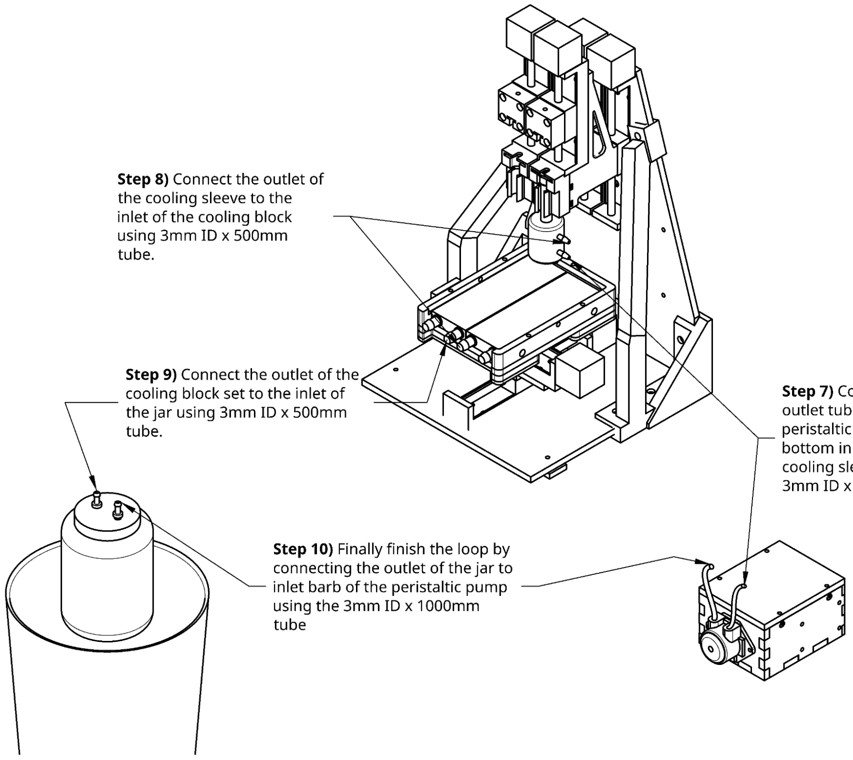

**Step 8)** Connect the outlet of the cooling sleeve to the inlet of the cooling block using 3mm ID x 500mm tube.

**Step 9)** Connect the outlet of the cooling block set to the inlet of the jar using 3mm ID x 500mm tube.

**Step 7)** Connect the outlet tube/barb of the peristaltic pump to the bottom inlet barb of the cooling sleeve using the 3mm ID x 250mm tube.

**Step 10)** Finally finish the loop by connecting the outlet of the jar to inlet barb of the peristaltic pump using the 3mm ID x 1000mm tube

**Tip-Tilt Platform Assembly**

Additional required components for tip-tilt platform

| Component | Quantity | Cost per unit | Total cost | Source of materials | Product/ Model number |
|---|---|---|---|---|---|
| Countersunk washer | 2 | $3.72 | $7.44 | McMaster | 92538A461 |
| 10 mm stainless steel ball | 1 | $1.443 | $1.443 | McMaster | 1598K32 |
| Extension spring | 2 | $3.298 | $6.596 | McMaster | 9433K196 |
| 2.5x12mm dowel pin | 4 | $0.2034 | $0.8136 | McMaster | 91585A293 |
| 3x20mm dowel pin | 2 | $0.3512 | $0.7024 | McMaster | 91585A390 |
| M3x0.5mm heat set insert | 2 | $0.2876 | $0.5752 | McMaster | 97171A310 |
| ¼”-80 fine-thread insert | 2 | $9 | $18 | McMaster | 98625A950 |
| ¼”-80 fine-thread thumb screw | 2 | $8.92 | $17.84 | McMaster | 97424A560 |
| Epoxy | 1 | $24.91 | $24.91 | McMaster | 7467A21 |

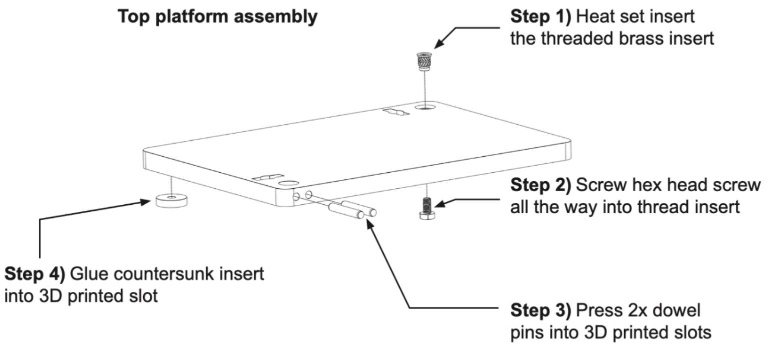

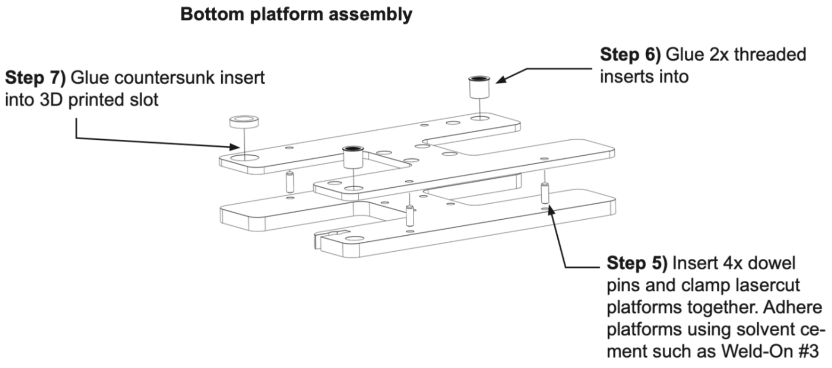

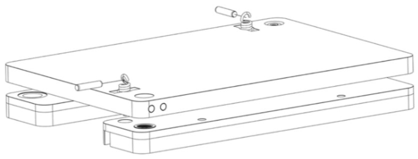

**Step 8)** Insert 2x springs through holes and lock in place by inserting dowel pins through the springs loop ends

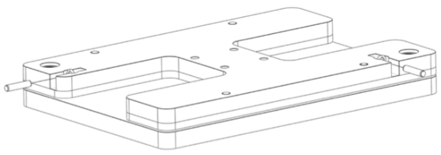

**Step 9)** Repeat step 8 on the opposite side, and make sure that the spring is properly attached on both sides

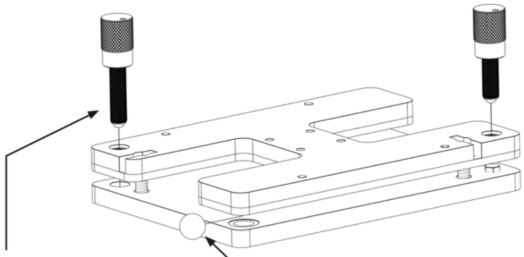

**Step 12)** Insert 2x screws to adjust platform to preferred tip and tilt

**Step 11)** Insert stainless steel ball between the countersunk washers
